# Fine-scale haplotype structure reveals strong signatures of positive selection in a recombining bacterial pathogen

**DOI:** 10.1101/634147

**Authors:** Brian Arnold, Mashaal Sohail, Crista Wadsworth, Jukka Corander, William P. Hanage, Shamil Sunyaev, Yonatan H. Grad

## Abstract

Identifying the forces that create and shape ecologically meaningful variation in bacteria remains an important challenge. For recombining bacteria, the sign and strength of linkage provide a unique lens into ongoing selection. We show derived alleles less than 300bp apart in *Neisseria gonorrhoeae* exhibit more coupling linkage than repulsion linkage, a pattern that cannot be explained by limited recombination or neutrality as these couplings are significantly stronger for nonsynonymous alleles compared to synonymous alleles. While linkage is shaped by many evolutionary processes, extensive simulations show only two distinct forms of positive selection can drive an excess of coupling linkage between neighboring nonsynonymous alleles: directional selection on introgressed alleles or selection that maintains distinct haplotypes in the presence of recombination. Our results establish a framework for identifying patterns of selection in fine-scale haplotype structure that indicate specific ecological processes in species that recombine with distantly related lineages or possess coexisting adaptive haplotypes.

## INTRODUCTION

Bacteria colonize diverse environments in which they experience challenges that leave distinct patterns in their genomes (Shapiro *et al.* 2009). As these environmental pressures are most often unknown, we rely on genomic signatures of selection to guide our understanding of bacterial ecology and evolution.

Unexpected patterns of linkage remain an important indicator of recent positive or negative selection, as many bacteria undergo frequent recombination (Vos and Didelot 2009) that unlinks loci. *N. gonorrhoeae* – a clinically important pathogen and an urgent public health concern due to growing incidence and antibiotic resistance – recombines extensively with closely- and distantly-related species and has been previously described as “panmictic” and “freely recombining” (O’Rourke and Stevens 1993; Smith *et al.* 1993). However, like in many bacterial species, proximate loci in *N. gonorrhoeae* exhibit stronger linkage than more distant loci (Arnold *et al.* 2018). This is as expected under neutral models that show neighboring polymorphisms are transferred together on recombination tracts, which range from tens to thousands of base pairs (Arnold *et al.* 2018; Lin and Kussell 2019). Many bacteria, including *N. gonorrhoeae*, also frequently exchange alleles with other species, thereby introducing clusters of linked mutations. While background levels of linkage are thus elevated for proximate loci, selection may also shape fine-scale haplotype structure, especially since the strength of selection acting on associations between neighboring loci may easily exceed the rate at which recombination breaks them. Consequently, methods to detect these signatures will aid understanding of microbial adaptation to ecological pressures and thus may inform the development of diagnostic and therapeutic measures to aid in control of bacterial pathogens.

Here, we develop an approach to study the interaction between selection and recombination within short distances. We compare the sign and strength of linkage between nonsynonymous (NSyn) derived alleles and synonymous (Syn) derived alleles, matched for relative recombination rate and allele frequency to control for population structure. Positive associations between derived alleles indicate coupling linkage, whereas negative associations indicate repulsion linkage. Using three large population genomic datasets of *N. gonorrhoeae*, we find all proximate loci exhibit an excess of coupling linkage, which cannot be explained by limited recombination. Intriguingly, NSyn alleles less than 300bp apart show significantly stronger couplings than Syn alleles. Many types of selection affect patterns of linkage, but extensive simulations show that only two specific forms of positive selection can create an excess of coupling linkage between neighboring NSyn alleles: adaptive interspecies admixture and balancing selection, defined here as spatially- or temporally-variable. We found this pattern is driven by a subset (∼10%) of exceptionally diverse genes, some of which exhibit evidence of interspecies recombination. Included among these genes are many metabolic proteins, but also membrane proteins and the Mtr operon that we previously showed contains admixed alleles that confer antibiotic resistance (Wadsworth *et al.* 2018).

Directional linkage measures applied to recombining bacteria that exchange alleles with other species can thus provide a wealth of information about selection within genes, information that may go undetected by linkage metrics that ignore sign or by conservative tests for selection that require more NSyn variation compared to Syn variation (*dN*/*dS* > 1).

## RESULTS

### Neighboring polymorphisms exhibit an excess of coupling linkage

Using three independent genomic datasets of clinical isolates collected from the United Kingdom (UK; n=214), New Zealand (NZ; n=148), and the United States (US; n=149), we tested the hypothesis that *N. gonorrhoeae* is a single recombining population under neutrality using several metrics to evaluate patterns of pairwise linkage. We first analyzed Syn single-nucleotide polymorphisms (SNPs) that, if neutral in effect, should provide a less biased view of recombination and population structure. To quantify linkage, we calculated the squared correlation coefficient *r*^2^, but we also computed the expected value of *D* (Methods) to measure the sign of linkage between alleles as it contains important information about haplotype structure. Positive values of E[*D*] indicate that alleles tend to co-occur (coupling phase), whereas negative values indicate that alleles tend to reside on different genetic backgrounds (repulsion phase). The sign of E[*D*] depends on the alleles one chooses to pair at two polymorphic loci, and we quantified *D* between derived alleles as in Takahasi and Innan (2008) by inferring the ancestral allele using one or several outgroups (with similar results; Methods). Derived alleles represent *de novo* mutations in *N. gonorrhoeae* or haplotypes imported from diverged populations that harbor alleles not found in the outgroups.

In a single recombining population under neutrality, E[*D*] is zero between allele pairs separated by any distance from equal amounts of coupling and repulsion linkage (Hill and Robertson 1968; Ohta and Kimura 1969, 1971). Less recombination between close alleles increases the variance of *D* (and thus values of *r*^2^), but these values of *D* take on positive or negative values with equal frequency such that E[*D*]=0 (Figure S1).

A variety of evolutionary processes may create coupling linkage, or positive E[*D*], including neutral dynamics such as population structure and interspecies admixture (Martin *et al.* 2006), or processes involving selection such as hitchhiking, balancing selection, or positive epistasis (antagonistic epistasis between deleterious mutations, or synergistic epistasis between beneficial mutations; Eshel and Feldman 1970; Thomson 1977; Charlesworth *et al.* 1997). Repulsion linkage, or negative E[*D*], is caused by clonal interference between mutations or negative epistasis (synergistic epistasis between deleterious mutations, or antagonistic epistasis between beneficial mutations; Hill and Robertson 1968; Sohail *et al.* 2017). If any of these evolutionary processes that create coupling or repulsion linkage are weak compared to the amount of recombination, E[*D*] will remain near zero as recombination rapidly breaks nonrandom associations.

In *N. gonorrhoeae*, we observed that linkage was generally low as expected due to extensive recombination, and *r*^2^ approached near-zero values as SNPs became separated by distances longer than ∼3kb (Figure 1A), in agreement with previous work (Arnold *et al.* 2018). However, for close SNPs that exhibited stronger linkage, the sign of E[*D*] became systematically positive (Figure 1B); neighboring Syn alleles were coupled more often than expected in a single neutral population, particularly for alleles separated by less than 300bp (Figure 1B).

**Figure 1.**
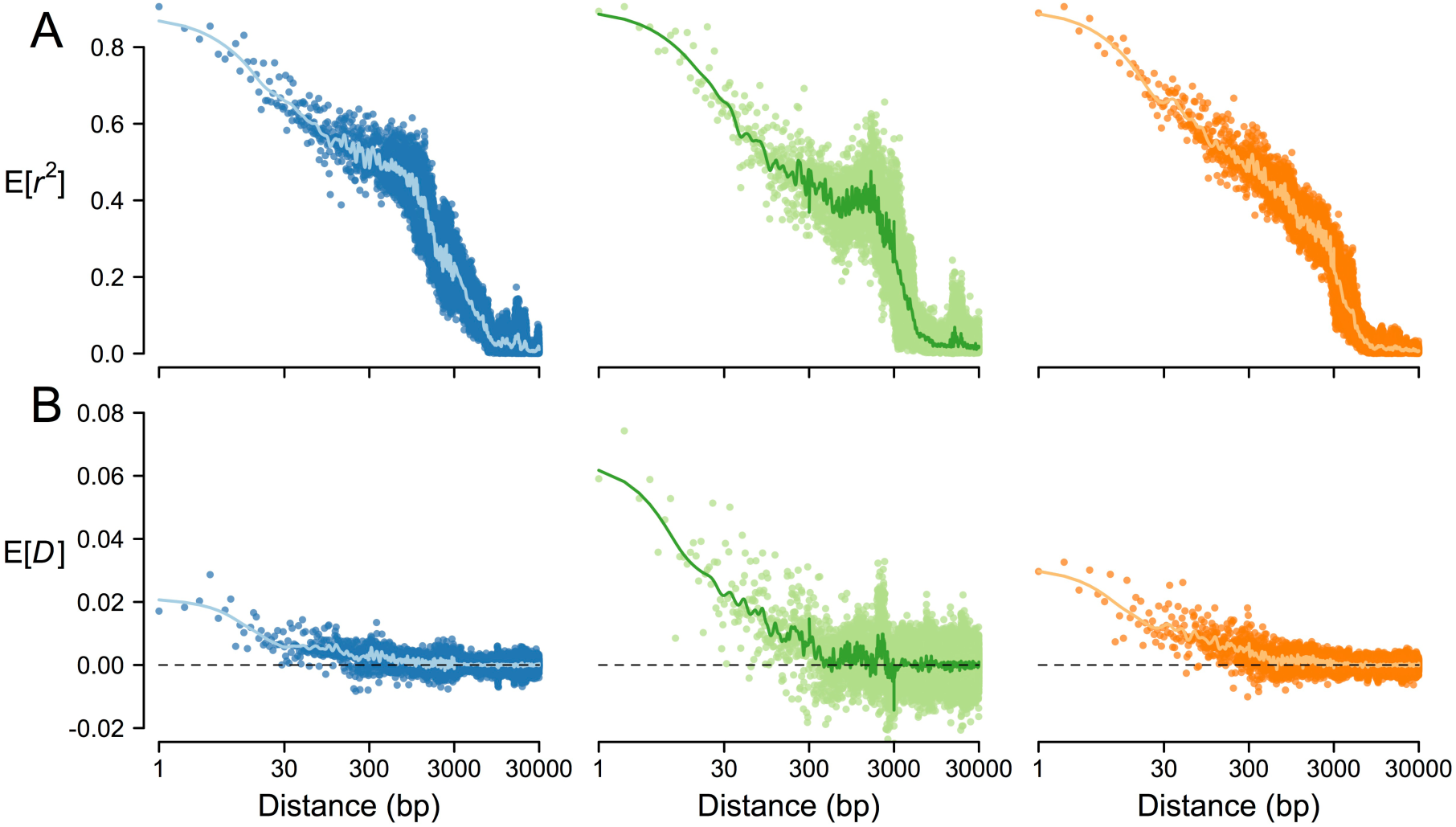
Patterns of linkage in *N. gonorrhoeae*. **(A)** Pairwise linkage between Syn SNPs, as measured by *r*^2^, reached very low levels for SNPs separated by >3kb. **(B)** However, when measured as *D*, pairwise linkage showed an excess of couplings between close SNPs that break down into random associations for distantly spaced SNPs >3kb apart. Each point represents the expected value of *r*^2^ or *D* for all SNP pairs separated by a number of base pairs shown in the x-axes. Results are shown for UK (blue), NZ (green), and US (orange).

### Selection shapes patterns of coupling linkage

Assuming NSyn SNPs are the actual targets of selection, we directly examined the role of selection in shaping this excess of short-range couplings by comparing the degree of coupling and repulsion linkage between Syn and NSyn alleles. Instead of using *D*, we quantified this linkage using the signed correlation coefficient *r* (separate from *r*^2^ in Figure 1A) that accounts for the dependency of *D* on allele frequencies (Hill and Robertson 1968; Methods) and – unlike the commonly used linkage metrics *r*^2^ and |*D*’| – preserves the sign of *D*.

Specifically, we evaluated whether *r* between NSyn derived alleles (*r*_N_) differed significantly from *r* between Syn derived alleles (*r*_S_). Because recombination breaks linkage and brings the sign of associations to zero, we controlled for potential differences in recombination between NSyn and Syn alleles in several ways to confirm that selection, not recombination, drives differences between *r*_N_ and *r*_S_. Since the probability that recombination unlinks two alleles varies with distance (Figure 1A), we first binned allele pairs by distance intervals of 100bp. If one allele category had more observations (e.g. more Syn than NSyn SNPs), we subsampled it 100 times to estimate variability. If NSyn and Syn alleles experienced similar selective pressures, *r*_N_ would be similar to *r*_S_ within each bin, but Figure 2 shows that for genomic distances less than ∼3kb, *r*_N_ was significantly different from *r*_S_. Intriguingly, for distances of less than ∼300bp where we observed the strongest coupling linkage for Syn alleles (Figure 1B), *r*_N_ was greater than *r*_S_, indicating NSyn alleles exhibit even stronger couplings (Figure 2A).

**Figure 2.**
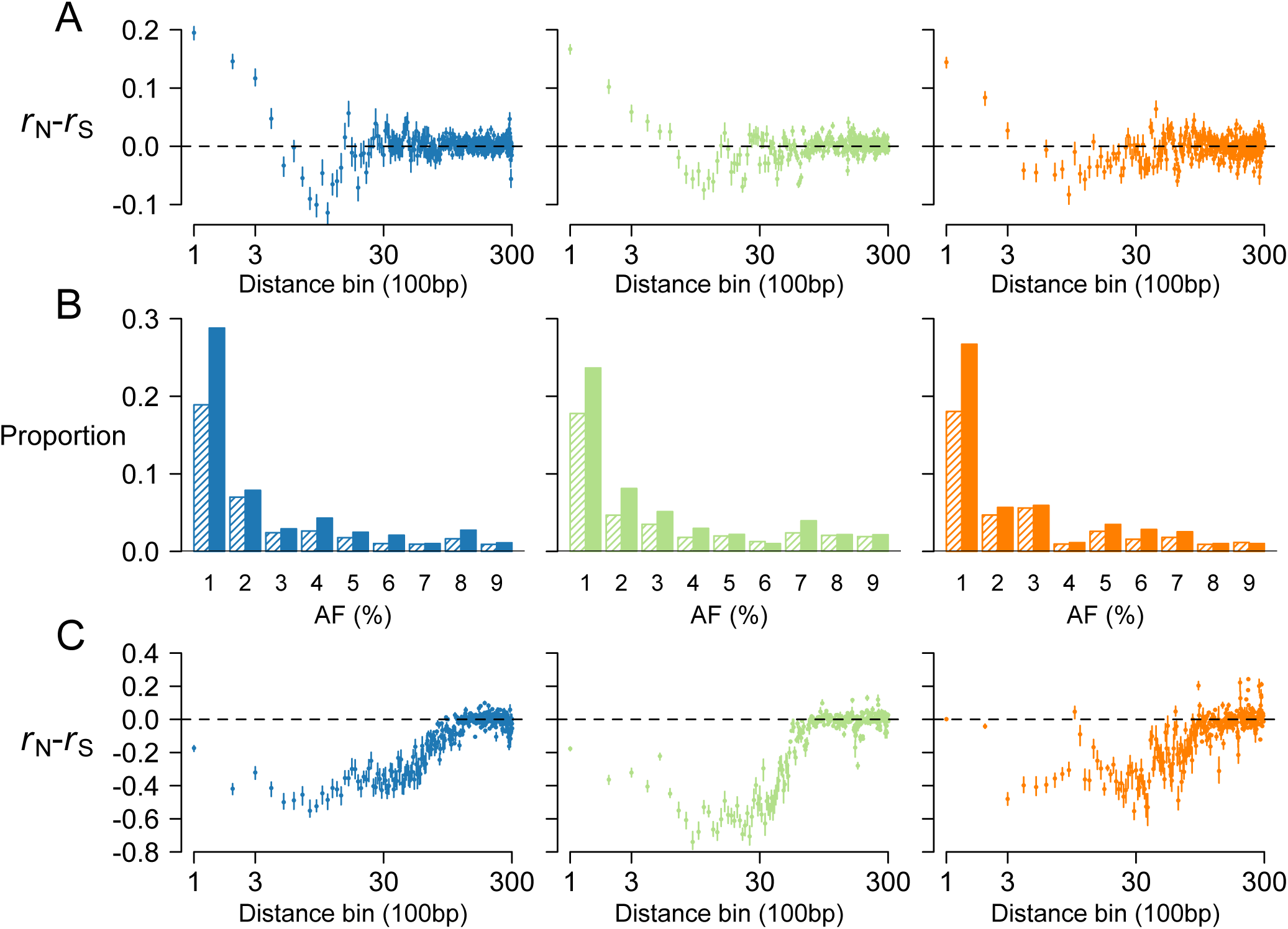
NSyn associations between close SNPs. **(A)** We binned SNP pairs by distance (100bp intervals) and compared NSyn *r* (*r*_N_) to Syn *r* (*r*_S_) for pairs within the same distance bin, downsampling the category that had fewer observations (*i.e.* NSyn or Syn) 100 times. **(B)** Allele frequency (AF) proportions for rare NSyn (solid) and Syn (shaded) SNPs. **(C)** *r*_N_-*r*_S_ calculated only for rare SNPs less 1% frequency in the sample. Dots **(A,C)** show the median value of resampled replicates, and the vertical lines connect the 5% and 95% quantiles. Results are shown for UK (blue), NZ (green), and US (orange).

We also controlled for potential differences in recombination between NSyn and Syn alleles by considering sample frequencies, which differed between allele categories (Figure 2B). Rare alleles are generally younger than those at higher frequencies and have had less opportunity for recombination to break down their linkage. Although we matched allele frequencies across the entire spectrum (below), we first considered rare alleles to study how negative selection shapes patterns of *r*–*r*, as rare polymorphisms are enriched for deleterious mutations. Negative selection is a pervasive evolutionary force in bacteria (Hughes 2005) that will skew NSyn alleles towards rare frequencies, make them younger than nearly-neutral alleles at the same frequency, and create clonal interference (or repulsion linkage) particularly when the strength of this selection exceeds the rate of recombination (Hill and Robertson 1968; Felsenstein 1974; McVean and Charlesworth 2000).

In *N. gonorrhoeae*, NSyn SNPs contained more rare alleles than Syn SNPs (Figure 2B). When we only considered alleles less than 1% frequency, which comprised ∼25% of all NSyn SNPs, we found *r*–*r* was significantly negative between neighboring SNPs – indicating an excess of NSyn repulsion linkage – but near zero between distant SNPs (Figure 2C). This is expected under a model of increasing clonal interference between deleterious mutations that experience less recombination, but negative selection with synergistic epistasis may also play a role (Eshel and Feldman 1970; Sohail *et al.* 2017). Nonetheless, since we observed an excess of repulsion linkage between rare NSyn alleles, negative selection is an unlikely explanation for the overall excess of coupling linkage between proximate NSyn alleles (Figure 2A).

### NSyn coupling linkage driven by positive selection

Focusing on short distances of ∼300bp, we found a significant excess of NSyn couplings, or positive *r*–*r*, for common alleles at 20-80% frequency that are nearly neutral or experiencing positive selection (Figure 3A; *P* ≤ 1.3×10^−7^ by Wilcoxon rank-sum test). We again controlled for potential differences in recombination between NSyn and Syn alleles by further binning these intermediate-frequency alleles into 10% frequency intervals, as higher frequency alleles (that may consist primarily of Syn mutations) are generally older and have had more time to experience recombination. Values of *r*–*r* for alleles within each interval were generally positive (Figure S2A), and meta-analysis across all intervals using Stouffer’s method indicated *r*_N_–*r*_S_ was significantly positive (*P* ≤ 5.9×10^−7^ for all three datasets; Table S1). Additionally, we further controlled for recombination by categorizing genes by their density of DNA Uptake Sequences (DUSs; Figure S2B), 12bp motifs that dramatically increase the uptake of exogenous DNA for homologous recombination (Goodman and Scocca 1988; Elkins *et al.* 1991; Ambur *et al.* 2007), such that genes bearing these motifs are significantly more likely to have their homologous alleles collected by the cell for recombination. Calculating *r*_N_–*r*_S_ only for genes that have low, medium, or high densities of DUSs showed similar results (Figure S2C), and meta-analysis across all DUS categories again showed *r*_N_–*r*_S_ was significantly positive (*P* ≤ 3.3×10^−10^ for all three datasets; Table S2). We also found similar results when we simultaneously controlled for both allele frequency and DUS density by binning alleles into 10% frequency intervals within each DUS density category (*P* ≤ 2.6×10^−4^; Table S3).

**Figure 3.**
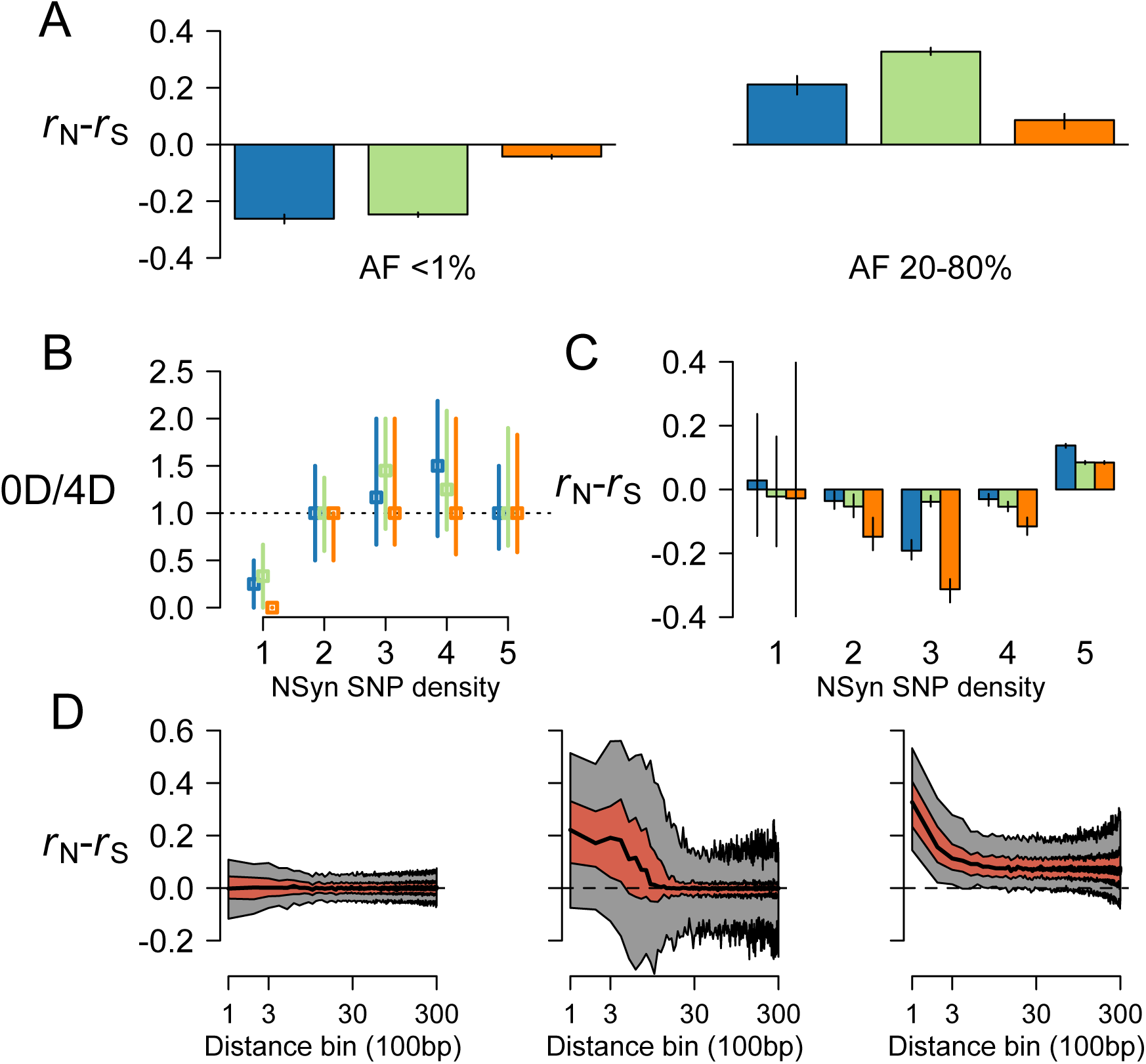
Greater NSyn associations for intermediate-frequency SNPs. **(A)** Focusing only on SNPs within 300bp, we calculated *r*_N_-*r*_S_ for SNPs with an allele frequency (AF) less than 1% (left) or between 20-80% (right). **(B)** The median ratio of diversity at zerofold and fourfold degenerate sites (OD and 4D, respectively; y-axis) for genes in higher NSyn SNP density quantiles was near one, and vertical lines indicate the interquartile range. **(C)** *r*_N_-*r*_S_ was significantly greater than zero only for genes with many NSyn SNPs. Results are shown for UK (blue), NZ (green), and US (orange). **(D)** We simulated interspecies recombination with beneficial mutations that have either nearly-neutral (*Ns*=0.1; left) or intermediate (*Ns*=25; middle) effect sizes. We also simulated spatially-variable selection with two demes (right) where mutations are beneficial in one deme but deleterious in the other (*Ns*=±4). The central black line represents the median *r*_N_-*r*_S_, the red region spans the interquartile range of medians, and the gray regions spans the 5% to 95% quantile of medians across 300 replicates.

A more precise approach to control for allele frequency differences would be to compare NSyn and Syn SNP pairs that have exactly matching frequencies, as opposed to binning alleles into 10% frequency intervals. While this approach discards a substantial number of allele pairs and may limit the ability to detect significant differences, we found exactly matching sets of NSyn and Syn common alleles within 300bp in our largest sample from the UK. We again found an excess of NSyn couplings (*r*_N_–*r*_S_=0.012, with values ranging from 0.0006 to 0.025 after subsampling).

To further explore whether NSyn coupling linkage is shaped by positive selection, we categorized core genes by NSyn SNP density, which reflects the dominant mode of selection: genes with fewer NSyn SNPs, which predominately experience negative selection, had less zero-fold degenerate (0D) diversity compared to fourfold-degenerate (4D) diversity, whereas genes with many NSyn SNPs, which likely experience positive selection, had 0D/4D ratios near or above one (Figure 3B). This is likely not due to Syn SNPs experiencing stronger negative selection in genes with high NSyn SNP densities, since Syn density also increased with NSyn density (Figure S3).

We found that the top 20% of genes that contained the most NSyn SNPs drive the overall pattern of positive *r*_N_-*r*_S_ between close alleles (Figure 3C). Furthermore, when we restricted the same analysis to variants below 5% sample frequency, which enriches NSyn alleles for those experiencing negative selection, we found *r*_N_-*r*_S_ was predominately negative (Figure S4). This agrees with the negative values of *r*_N_-*r*_S_ in Figure 2C for alleles less than 1% frequency and again demonstrates that negative selection does not drive the observed excess of NSyn couplings within genes.

The fact that genes with an excess of NSyn couplings also had exceptionally high diversity suggests balancing selection may be involved, as this increases local SNP densities by maintaining haplotypes at selected and linked loci. While directional selection typically eliminates diversity, we may observe local increases in SNP density if it is currently spreading introgressed alleles. Indeed, adaptive interspecies admixture and balancing selection affected *r*_N_-*r*_S_ in similar ways according to simulations (Figure 3D). For simulations of admixture with neutral effects, *r*_N_-*r*_S_ for common alleles within 300bp was significantly positive (α=0.05) for 5.1% of replicates, compared to 41% and 100% of replicates for simulations of adaptive admixture and balancing selection, respectively.

We also calculated *r*_N_-*r*_S_ from simulations of positive and negative directional selection in a single population, with and without positive epistasis. None of these scenarios created a significant excess of NSyn couplings for proximate alleles (see Supplementary Results for more information about simulations). Thus, accounting for interspecies admixture may provide additional insight into the mechanisms underlying NSyn coupling linkage, especially considering *N. gonorrhoeae* is known to recombine with closely related species (Spratt *et al.* 1992; Feil *et al.* 1996; Hanage *et al.* 2005; Corander *et al.* 2012; Ezewudo *et al.* 2015; Wadsworth et al. 2018).

### NSyn coupling linkage associated with interspecies admixture

We assembled genomic datasets for three closely related species: *N. meningitidis, N. polysaccharea*, and *N. lactamica* (see methods section) to characterize genome-wide allele-sharing between species. A Neighbor-Joining tree of all four species agreed with the evolutionary relationships described in previous studies (Figure 4A; Bennett *et al.* 2013, 2014). Figure 4B shows intragenic diversity in *N. gonorrhoeae* was highly variable along the genome, with regions of high SNP density punctuated by regions containing few SNPs. When we compared *N. gonorrhoeae* with *N. meningitidis* (Figure 4B), peaks in SNP density visually corresponded with more shared polymorphism and less monophyletic genealogies, as measured by the genealogical sorting index (gsi), which ranges from 0 (completely mixed) to 1 (reciprocal monophyly). Indeed, when we binned genes by NSyn SNP density, those with higher densities tended to have more shared polymorphism with *N. meningitidis* and lower values of gsi (Figure 4C). We observed similar patterns when comparing *N. gonorrhoeae* to *N. polysaccharea* and *N. lactamica*, although as expected, the range of gsi values became more skewed towards 1 as genetic divergence between species increased (Figure 4B, Figure S5).

**Figure 4.**
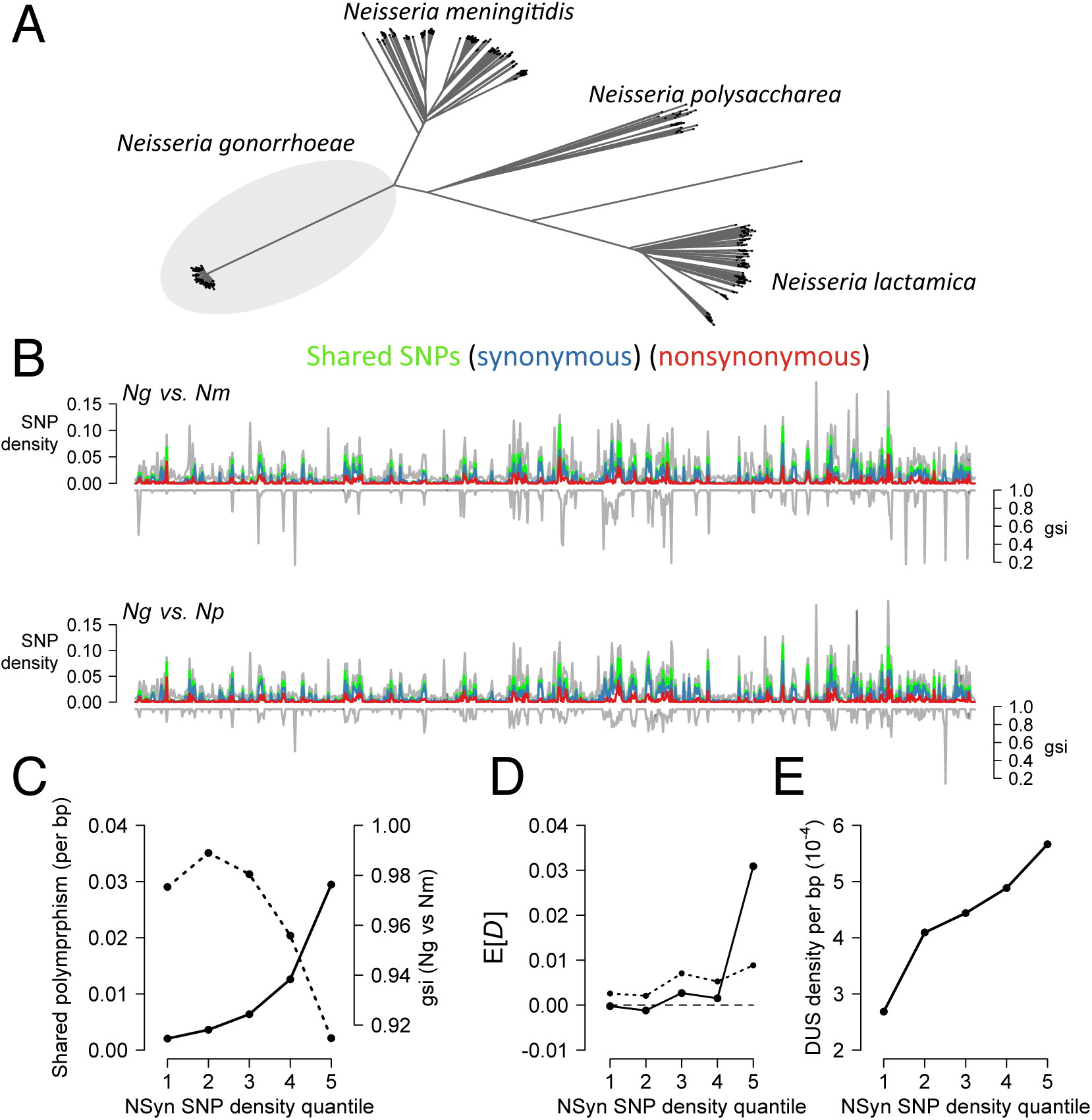
Gene diversity correlated with interspecific-shared polymorphism and lower gsi values. (**A**) Unrooted Neighbor-Joining tree constructed from pairwise distances for three *Neisseria* species: *N. gonorrhoeae* (*Ng*), *N. meningitidis* (*Nm*), and *N. polysaccharea* (*Np*). (**B**) Diversity in *Ng* (only UK dataset shown), quantified as SNP density (gray lines in top panels), for each gene showed those with many SNPs also share many of these polymorphisms (green lines) with *Nm* (upper) or *Np* (lower). Blue and red lines represent shared Syn and NSyn SNPs, respectively. Genes with many SNPs also had lower values of gsi. **(C)** Categorizing genes by NSyn SNP density showed those with more diversity have more shared polymorphism (solid line) and lower values of gsi (dotted line). Genes with higher NSyn SNP densities also had higher mean values of *D* **(D)**, as measured between Syn SNPs (dotted line) or NSyn SNPs (solid line), and more DUSs **(E)**. In **(B)**, genes are ordered in these plots according to their relative position in the FA1090 reference genome for *Ng*.

While interspecies recombination explains these trends, incomplete lineage sorting (ILS) may also contribute to shared ancestry, especially considering *N. meningitidis* and *N. gonorrhoeae* are sister species. However, while admixture between diverged populations may create coupling linkage, as we observed in *N. gonorrhoeae* (Figure 4D), we show through simulations that ILS does not produce this pattern (Figure S6). Genes with more NSyn SNPs also tended to have more DUSs (Figure 4E), which are known to significantly enhance transformation, and all *Neisseria* species analyzed here share the same DUS (Frye *et al.* 2013).

In summary, genes with a high density of NSyn SNPs not only had a significant excess of NSyn coupling linkage (Figure 3C) but also displayed evidence of interspecies admixture (Figure 4). An excess of NSyn couplings was also directly related to interspecies admixture, as individual genes in *N. gonorrhoeae* in which *r*_N_-*r*_S_ > 0.2 also had lower values of gsi and more shared polymorphism with the three close relatives included here (Figure S7), as well as slightly higher levels of intragenic DUSs (0.00059/bp vs. mean of 0.00044 for all genes). While gsi uses gene phylogenies to measure shared ancestry, a complementary, higher resolution analysis using fastGEAR (Mostowy *et al.* 2017) also showed how ancestry changes across the length of admixed genes, revealing tracts of DNA from other species or entire alleles that have no identifiable DNA from *N. gonorrhoeae* (Figure S8). Collectively, these observations highlight the potential role of interspecies admixture in driving NSyn couplings within highly diverse genes under positive selection.

### Outlier genes with an excess of NSyn coupling linkage

The distribution of *r*_N_-*r*_S_ calculated for individual genes was multimodal for two datasets, with most genes having near zero values (Figure 5). Although *r*_N_-*r*_S_ outlier genes generally had more NSyn polymorphism, of the 29 outlier genes we detected in at least one dataset (*r*_N_-*r*_S_ > 0.2, Methods), only one exhibited *dN*/*dS* > 1 (Table S4). Twenty of these genes had annotation information and over half (14) were involved in metabolic processes (Table S4), suggesting selection on metabolism creates an important component of haplotype structure in *Neisseria*. Of the other outliers, two were membrane proteins, and another was *mtrE*, a candidate for adaptive admixture, as it is part of an operon encoding an efflux pump that has acquired antibiotic resistance via recombination with several closely related species (Wadsworth *et al.* 2018). Many of these *r*_N_-*r*_S_ outlier genes also had a large excess of intermediate-frequency variants (Table S4), a pattern expected under balancing selection. Of these, one encodes a membrane protein, a category that is frequently documented to experience such selective pressures (e.g. Gupta *et al.* 1996), and two others encode metabolic proteins that act in different parts of the same amino sugar and nucleotide sugar pathway. These balancing selection candidates also did not exhibit evidence of interspecies admixture according to gsi values (at least for the three outgroup species used here), such that an excess of NSyn couplings may be driven solely by balancing selection on intraspecific diversity.

**Figure 5.**
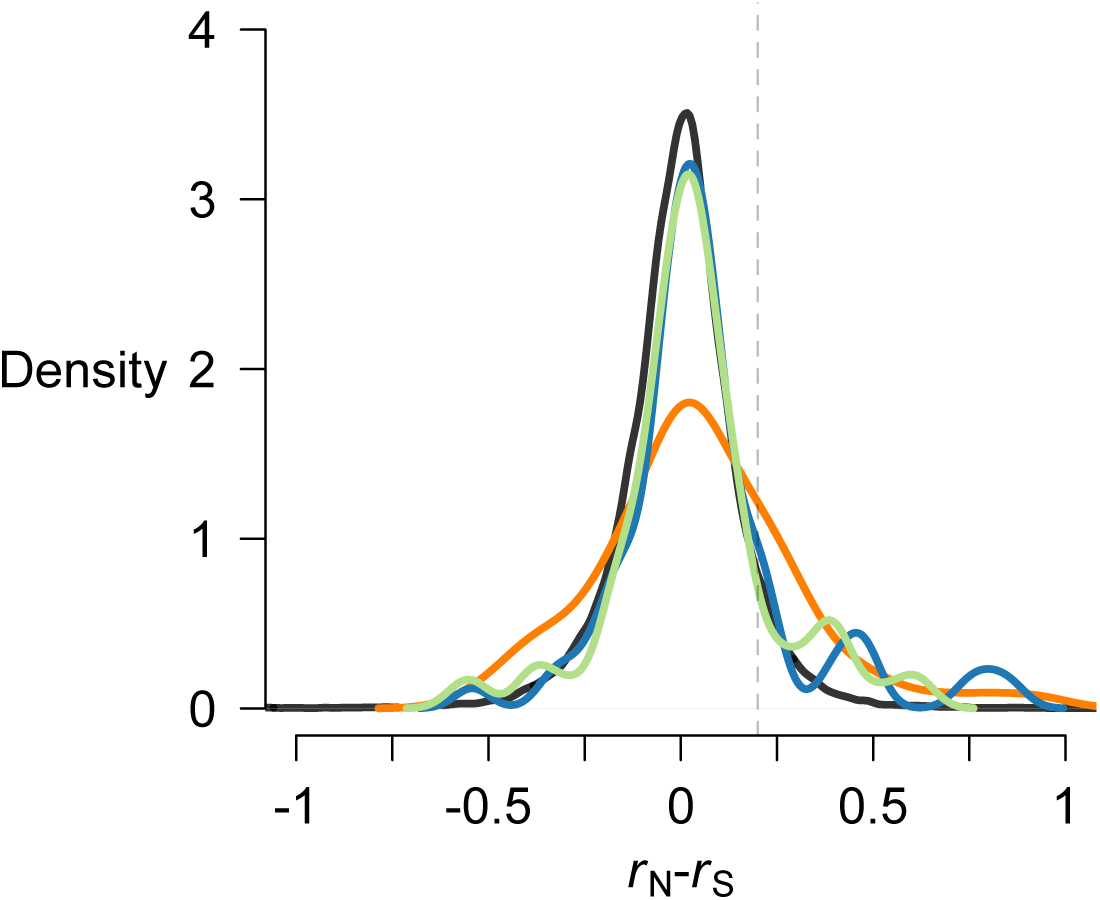
Distribution of intragenic *r*_N_-*r*_S_ values. Values of *r*_N_-*r*_S_ by gene showed a multimodal distribution with an excess of NSyn couplings compared to neutral simulations (black). The dashed gray line indicates the threshold used for *r*_N_-*r*_S_ outliers. For each gene with at least 5 Syn and 5 NSyn SNPs, *r*_N_-*r*_S_ was calculated for derived alleles with frequencies above 5%, Results are shown for UK (blue), NZ (green), and US (orange).

## DISCUSSION

*N. gonorrhoeae* is a highly recombining human pathogen that has stably colonized multiple ecological niches, including the urogenital tract, the oropharynx, and the rectum, and has evolved resistance to all clinically important antibiotics used to treat infection. Given the pervasiveness of recombination within and between *Neisserial* species, analyses that jointly study selection and recombination could reveal novel insights into the biology of the *Neisseria* that other approaches will not detect. Using patterns of directional linkage between derived mutations in *N. gonorrhoeae*, we show associations between neighboring polymorphisms are shaped by a form of selection that creates stronger coupling linkage between NSyn alleles compared to Syn alleles (*r*_N_-*r*_S_>0). Other summary statistics show at least some of this positive selection may act on alleles introgressed from closely related species, and simulations with adaptive admixture between diverged populations or balancing selection produce similar patterns.

These two forms of selection, which are not mutually exclusive, inflate NSyn couplings between close alleles in different ways. In the case of balancing selection, which maintains allelic variation in a large panmictic population that encounters heterogeneous environments, NSyn couplings represent allelic combinations that are beneficial in one environment but neutral or deleterious in others. While recombination occurs between all lineages and breaks linkage, selection maintains preferred allele combinations if it is sufficiently stronger than recombination, a scenario that becomes increasingly likely for neighboring alleles that experience less recombination (Wiuf and Hein 2000; Lin and Kussell 2019). In the case of adaptive admixture, linked beneficial NSyn alleles are introduced on short recombination tracts that rapidly increase in frequency, which makes them younger (providing less opportunity for recombination to break their linkage) than Syn alleles that rose to the same frequency via drift. This contrasts starkly with beneficial mutations that randomly arise within a species, as these typically display repulsion linkage from clonal interference due to limited recombination (Figure S18, Hill and Robertson 1968). Importantly, simulations of interspecies recombination involving neutral mutations do not tend to produce positive values of *r*_N_-*r*_S_ (Figure 3D).

The *r*_N_-*r*_S_ metric developed here serves as a complementary way to study selection in bacterial and other sexual organisms, as it detects signatures of selection that differ from those identified by other methods, such as *dN/dS* or metrics that compare diversity with divergence (McDonald and Kreitman 1991; Stoletzki and Eyre-Walker 2011). Intriguingly, an excess of couplings between close Syn alleles within ∼300bp, as in Figure 1B, has also been documented in a highly recombining, thermophilic *Synechococcus* species (but not for NSyn alleles, as they were not studied; Rosen *et al.* 2015), and the authors hypothesize these associations reflect hitchhiking from spatially-variable selection maintaining linked mutations. These findings also have important implications for estimating recombination parameters from bacterial genomic data using neutral models (e.g. Lin and Kussell 2019), as selection may shape short-range patterns of linkage.

Although numerous unknown selective pressures may be driving the maintenance of diversity in *N. gonorrhoeae*, epidemiological data provide promising clues. *N. gonorrhoeae* thrives within different sexual networks in which lineages are transmitted through diverse habitats that may present distinct challenges. For instance, in networks involving men who have sex with men, *N. gonorrhoeae* has been isolated from the urethra, rectum, and oropharynx, and for networks involving women, its niches include the vagina/cervix (Grad *et al.* 2014; De Silva *et al.* 2016; Lee *et al.* 2017; Sánchez-Busó *et al.* 2018). Adaptive admixture and the coexistence of diversity could thus reflect spatially- or temporally-variable selection from lineages sojourning in these distinct niches.

Introgressed diversity could also represent recently acquired alleles that are unconditionally beneficial and currently spreading through the entire species. *N. gonorrhoeae* has substantially less diversity than all of its close relatives (Figure 4A), which may reflect a genetic bottleneck that accompanied recent speciation, such as a single *N. meningitidis* lineage giving rise to the present-day species. Severe bottlenecks may fix rare, deleterious variation present in the ancestral population, such that adaptive admixture may simply reflect the acquisition of beneficial wild-type alleles (e.g. Kim *et al.* 2018).

Additional analyses offer the opportunity for a more complete understanding of genome-wide evolution in *N. gonorrhoeae*. Here, we only consider core genes and focus on contemporaneous samples by subsampling larger datasets to obtain isolates from a restricted time period (e.g. a single year). Analyses that include longitudinal samples will help understand the temporal dynamics of mutations and may be particularly useful for establishing types of positive selection, for instance whether directional selection on introgressed haplotypes or balancing selection shapes patterns of variation at a specific locus, as the latter scenario will maintain polymorphisms over time. Moreover, a deeper understanding of the other commensal species within *Neisseria*, such as any species *N. gonorrhoeae* may recombine with, will shed additional light on *N. gonorrhoeae* biology and speciation processes within the genus.

## METHODS

### Sequence data processing

We reanalyzed three previously published sequencing datasets of *N. gonorrhoeae* isolates collected from Brighton (United Kingdom; De Silva *et al.* 2016), New Zealand (Lee *et al.* 2017), and the United States (Grad *et al.* 2014, 2016). Since sequencing errors may give rise to structure in genomic datasets (and thus positive E[*D*]) if subsets of isolates are more likely to have erroneous nucleotides, we took many precautions to ensure high-quality DNA alignments for downstream analysis.

We analyzed the quality of raw reads using FastQC (Andrews 2010), and samples with GC content that differed more than 2.5 standard deviations from the mean were not included in analyses (3 isolates in the dataset from Brighton). These reads were used to create *de novo* assemblies using SPAdes (v. 3.11; Bankevich *et al.* 2012), and contigs were joined into larger scaffolds using SSPACE (Boetzer *et al.* 2011). We mapped raw reads back to these assemblies using SMALT (v. 0.7.6), using only those reads in which at least 95% of bases successfully mapped. With these mapped alignments, we used Pilon (v. 1.13; Walker *et al.* 2014) to correct any nucleotides not supported by the raw read data. For a maximum of four iterations, we repeated this process of mapping reads back to the *de novo* assembly and correcting nucleotides, unless Pilon made no changes to the updated assembly.

Using these polished assemblies along with information from the final mapped alignment, we marked nucleotide positions as “N” if they met any of the following criteria: (1) alignment depth less than 5 bases, (2) mean base quality less than 20, (3) mean mapping quality less than 20, (4) alignment depth more than 1.8 times the median depth across all positions, (5) positions flagged as “Amb” by Pilon that have significant evidence of more than one allele, or (6) positions in which a second allele was present with a frequency greater than 10% of all base calls. These positions marked as “N” were excluded from downstream analyses. In addition, we discarded all contigs that were less than 300 bp or had more than 50% of their sites masked by the filtering step above (primarily plasmids with very high sequencing depth).

### Gene annotation and alignment

We annotated assemblies with Prokka (Seemann 2014) using the proteome of the FA1090 reference genome. We then used Roary (Page *et al.* 2015) to identify core genes present in all *N. gonorrhoeae* isolates, defining orthologous genes as having at least 90% amino acid similarity. This threshold tends to misclassify core genes with very diverged alleles as multiple accessory genes (Ding *et al.* 2018), such that we may miss those that have admixed with distant species. However, we preferred a conservative set of core genes for accurate estimates of linkage, since erroneous clutstering of alleles may give rise to artifactual associations.

We then realigned these core genes with PAGAN (Löytynoja *et al.* 2012), a phylogeny-aware multiple sequence aligner. PAGAN performs an amino acid alignment that helps maintain the reading frame of diverged sequences, which is required for accurately labeling polymorphisms as Syn or NSyn. All position information between polymorphic sites was derived from the relative positions of genes in the FA1090 reference genome used to annotate *de novo* assemblies, not from a reference-based DNA alignment. While rearrangements may occur within *N. gonorrhoeae*, synteny between genomes likely extends beyond 3kb (Figure S9), the distance around which *r*^2^ and *D* approach near-zero values. Moreover, codons in the FA1090 reference sequence were used to ascertain the functional effect of polymorphisms, *i.e.* whether they were Syn or NSyn. For all downstream analyses, we excluded COGS that were present in multiple copies in at least one individual and all alignments that contained multiple premature stop codon polymorphisms.

### Polarizing mutations

For analyses that required polarized mutations (e.g. calculating *D*), we used progressiveMauve (Darling *et al.* 2010) to align the FA1090 reference genome to an *N. meningitidis* outgroup sequence to infer the derived and ancestral state of each bi-allelic polymorphism within *N. gonorrhoeae*. All analyses were done using the α14 *N. meningitidis* reference as the outgroup sequence, as it is the closest known reference to *N. gonorrhoeae* (Budroni *et al.* 2011). However, since this reference sequence may have acquired derived mutations since its divergence from *N. gonorrhoeae*, we also polarized mutations using one *N. meningitidis* (α14) and one *N. polysaccharea* (SRA accession number ERR976854) sequence. We only considered positions in which both outgroup sequences had the same nucleotide and *N. gonorrhoeae* had a biallelic polymorphism, with one of the alleles also found in the outgroup. The outgroup consensus allele served as the ancestral state. This method of polarization gave highly similar results (Figures S10). We also note that we included the FA1090 reference sequence when clustering COGs with Roary and realigning with PAGAN (above), so that we could map positions within gene alignments to those in the reference used in the multispecies Mauve alignment used to polarize mutations. However, this reference sequence was excluded from analyses of diversity and linkage within each of the three datasets.

### Data filtration

With these core genome alignments, we calculated the number of pairwise SNP differences between alignments using Disty McMatrixface (https://github.com/c2-d2/disty) and down sampled closely related isolates in order to exclude those from the same transmission chain. Clusters of closely related genomes could arise from reporting bias within contact networks, as isolates were sampled from symptomatic individuals that visited sexual health clinics. Specifically, we down sampled clusters of isolates that were identical (0 SNP differences) or separated by less than 6 SNPs to a randomly selected isolate within that cluster. We used the 6-SNP threshold because De Silva *et al.* (2016) showed that isolates collected from known contact networks in low-transmission settings had fewer than 6 SNP differences. For either SNP threshold, patterns of pairwise *D* looked highly similar (Figure S11), and all analyses shown here were performed on alignments down sampled according to the 6-SNP threshold. We also excluded isolates that were collected from the same patient over time (UK dataset), although these samples would have also been excluded using the SNP thresholds above.

Lastly, to avoid any potential temporal structure that may inflate E[*D*], we analyzed only isolates collected in 2013 from the UK data, and only those collected in 2010-2011 from the US dataset. All isolates in the NZ dataset collected from 2014-2015 were analyzed. Overall, we analyzed 214 sequences from UK, 149 from US, and 148 from NZ.

### Linkage statistics

We primarily measured linkage using *r*, where 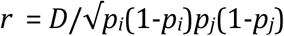 and *D* = *p*_*ij*_*-p*_*i*_*p*_*j*_, or the probability derived mutations at site *i* and *j* occur together minus the random expectation based on their individual frequencies (Hill and Robertson 1968). While *r* and *r*^2^ are commonly used to quantify linkage, their maximum values depend on the degree of symmetry between the allele frequencies of a pair of loci under consideration (VanLiere and Rosenberg 2008). However, the range of values of another linkage statistic, *D’* (Lewontin 1964), is independent of allele frequencies (Hedrick 1987). We repeated our *r*_N_-*r*_S_ analyses (Figure 2,3C) using *D’* and found qualitatively similar results: proximate derived NSyn alleles had more coupling linkage than derived Syn alleles separated by similar distances, and this observation was driven by genes with the highest density of NSyn SNPs (Figure S12).

We also note that for analyses in Figure 4D, while theory has shown that E[*D*]=0 for a panmictic species (Hill and Robertson 1968; Ohta and Kimura 1969, 1971), we confirmed with simulations that conditioning on SNP density does not alter this expectation (E[*D*|SNP density] ∼ 0; Figure S13), such that genes with many SNPs are not expected to have higher *D*, as we observe.

All data for *N. gonorrhoeae*, including gene alignments and input files from other software, and documented Perl scripts used in these analyses will be freely available on github (https://github.com/brian-arnold/NgonorrhoeaeLinkageGenomics) at the time of publication.

### Significance tests

When accounting for allele frequencies or DUS densities (Figure S2, Table S1, S2, S3), we tested if *r*_N_-*r*_S_ was significantly positive within each bin or category using the Wilcoxon rank-sum test (“Stats” package in R). To meta-analyze *P* values across bins, we used Stouffers Z-score method (“metap” package in R; Liptak 1958). We also applied these tests to simulated data (Supplementary Methods).

### Interspecies analysis

To study shared ancestry between *N. gonorrhoeae* and its close relatives, we downloaded all assemblies from NCBI or raw read data from the Short Read Archive (SRA) for *N. meningitidis, N. polysaccharea*, and *N. lactamica* on October 19, 2017. We created *de novo* assemblies from these raw reads using SPAdes as above. We then only used assemblies that had N50 greater than 10kb, no more than 150 contigs, and at least one contig that was at least 300kb. We also excluded several assemblies that were very diverged from the majority of samples for that species according to a visualization of Neighbor-Joining trees (1 for *N. meningitidis*, 15 for *N. lactamica*, and 4 for *N. polysaccharea*); these highly diverged isolates may have bad assemblies or were incorrectly labeled. In total we used 431 *N. meningitidis*, 326 *N. lactamica*, and 37 *N. polysaccharea* assemblies, and the accession numbers for these may be found in Table S5.

We then mapped sequences from each outgroup species to the *N. gonorrhoeae* FA1090 reference genome. However, instead of mapping raw reads, we first used them to create *de novo* assemblies (using SPAdes, as above) for each isolate and then mapped scaffolds from these assemblies to the reference using progressiveMauve (Darling *et al.* 2010). We opted for this approach since long scaffolds have more information about mapping position than short sequencing reads, and microsynteny among the four species extends beyond the length of a gene (Figure S9). For each gene we previously identified as “core” within the *N. gonorrhoeae* datasets, we extracted the sequences from these progressiveMauve alignments and incorporated them into existing core gene alignments with MAFFT (using the -add and -keeplength options; Katoh and Frith 2012). We excluded genes that were present in fewer than 20% of isolates in the outgroup species or with alignments in which over 50% of positions were gaps. While these alignments were directly used to quantify shared ancestry in terms of shared SNP density, we also constructed a multispecies phylogeny for each gene using RaxML (v. 8.1.5; Stamatakis 2014) with 20 bootstrap replicates under the GTRCAT model of rate heterogeneity. These phylogenies were used to calculate gsi (Cummings *et al.* 2008) with the genealogicalSorting R package, and we took the mean gsi value across all 20 bootstrap replicates for leaves labeled as *N. gonorrhoeae*.

### Forward-time simulations

To simulate bacterial evolution, we used *fwdpp* (Thornton 2014), a library of C++ functions that abstracts the essential tasks of forward-time simulations such that one may use them to design custom simulators. We modified the code to accommodate haploid populations and also made custom functions to simulate homologous recombination (*i.e.* gene conversion) and various types of selection. With *fwdpp,* we simulate Wright-Fisher metapopulations under an infinite-sites mutation model. For more information about details of simulation parameters and results, and the calculation of individual fitness, please see Supplementary Methods. Source code will be available on github at the time of publication.

### DUS identification and density estimation

To locate DUSs within *Neisseria* genomes, we searched sequences for the degenerate 10bp motif 5’-GCCGTCTGAA-3’, allowing one nucleotide to vary within the first three or last two positions since mutations within positions 4 to 8 may result in highly reduced rates of uptake (Frye *et al.* 2013). We searched both forward and reverse strands. When counting DUS densities within genes or their flanking regions (200bp), we collapsed DUSs within 300bp into a single observation: although increasing the number of DUSs generally increases DNA uptake and transformation, those separated by short distances exhibit interference (Ambur *et al.* 2012). DUS density per gene (including flanks) is highly variable (Figure S2B). When controlling for relative DUS density while calculating *r*_N_-*r*_S_, we created three categories (high, medium, and low) but excluded genes with either no DUSs or very high DUS densities (top 15%) to avoid categories with highly variable densities (Figure S2B).

### *dN/dS* analysis

Using *omegaMap* (Wilson and McVean 2006), we calculated *dN/dS* for genes that also had high values of *r*_N_-*r*_S_ (Table S4). For information about the parameters used in this analysis, please see the Supplementary methods.

## ACKNOWLEDGEMENTS

We would like to thank Kevin Thornton for his generous help with *fwdpp*. BJA was supported by a postdoctoral fellowship1 F32 GM120839-01, CBW and YHG were supported by the Richard and Susan Smith Family Foundation and National Institutes of Health grant R01 AI132606. JC was funded by the ERC grant 742158, and WPH was funded by National Institutes of Health grant U54 GM088558. The funders had no role in study design, data collection and analysis, decision to publish, or preparation of the manuscript. The computations in this paper were run on the Odyssey cluster supported by the FAS Division of Science, Research Computing Group at Harvard University.

## COMPETING INTERESTS

The authors declare no competing interests.

