## Supplementary Material for "Fine-scale haplotype structure reveals strong signatures of positive selection in a recombining bacterial pathogen"

*Figure S18. Simulations of panmixia with negative or positive directional selection and epistatic fitness effects.* 23

### SUPPLEMENTARY RESULTS

#### FORWARD-TIME SIMULATIONS TO DEFINE BEHAVIOR OF $R_N$-$R_S$.

##### Admixture with positive selection.

We designed a forward-time simulator to test whether selection on introgressed alleles can create the excess of NSyn couplings we observe at neighboring loci. To do so, we extended *fwdpp* (Thornton 2014) to model haploid populations with gene conversion, mimicking homologous recombination in bacteria.

Using *fwdpp*, we studied the simple scenario of a sink (or recipient) species receiving short segments of DNA from a diverged source (or donor) species. We modeled interspecies admixture by first evolving the donor and recipient populations in allopatry, allowing many neutral and beneficial mutations to fix in the donor species (Figure S14). After allopatric evolution, we specified a low chance (~2%) that an individual from the donor population serves as DNA template for recombination in the recipient population. This simulates the uptake of potentially many linked mutations, several of which may be beneficial, and mimics a biological scenario in which *N. gonorrhoeae* encounters exogenous DNA of close relatives, all of which may be occasionally found in the oropharynx. Sometime after the onset of admixture (e.g., $0.1N$ generations, where $N$ is the population size), we took a sample of size $n$=200 from the recipient population and analyzed the sequence data as in Figure 2, treating beneficial mutations as NSyn and neutral mutations as Syn.

If we took a sample from the recipient population soon after the onset of admixture (~$0.1N$ generations), $r_N$ had no tendency to be greater than $r_S$ when beneficial mutations from the donor species had nearly neutral effects in the recipient ($Ns < 1$; Figure S14B). However, $r_N$ became systematically greater across short genomic distances, but not longer ones, as beneficial effect sizes became intermediate in strength ($Ns$=25; Figure S14C, Figure 3D). These were the largest beneficial effect sizes we simulated for adaptive admixture.

For these simulations of adaptive admixture with $Ns$=25, we tested whether $r_N$ was significantly greater than $r_S$ using the Wilcoxon rank-sum test in two ways. We first applied this test to all common alleles (20-80% frequency) within 300bp. We found significantly greater coupling linkage between neighboring NSyn alleles in 41% of simulation replicates, at the $\alpha$=0.05 significance level. We then further controlled for allele frequency differences between Syn and NSyn SNPs by binning common alleles into 10% frequency intervals, performing a Wilcoxon rank-sum test for each interval, and meta-analyzing all $P$ values using Stouffers Z-score method. This analysis indicated that only 9% of simulation replicates had significantly more NSyn coupling linkage. The differences between these analyses may be driven by a loss of power from fewer observations of $r_N$ and $r_S$ after binning common alleles into 10% frequency intervals. We note that for simulations of balancing selection (below), we performed similar statistical tests and observed much smaller differences between analyses that used values of $r_N$ and $r_S$ calculated for all common alleles and those that used values calculated from common alleles binned into 10% frequency intervals.

We also explored varying selection intensities and took samples at different evolutionary times after the onset of admixture (Figure S15), showing the signal of $r_N$-$r_S$ >0 arose soon after admixture starts (~0.02$N$ generations) and persisted for relatively long periods of time (until ~2$N$ generations, with continual admixture). While the exact dynamics over evolutionary time depend on the rate of admixture and the number of diverged donor populations available, these simulations show that simple models of adaptive admixture involving short fragments of donor DNA containing beneficial mutations can significantly inflate NSyn coupling linkage over short genomic intervals. Neither epistasis between mutations nor balancing selection is required.

Simulations also indicated $r_N$-$r_S$ may be negative for SNPs separated by distances just longer than the mean admixture tract length (Figure S14C, S15), which we modeled as ~300bp since $r_N$-$r_S$ >0 across this genomic interval (Figure 2). This likely represents clonal interference or Hill-Robertson interference (Hill and Robertson 1968) between independent, nearby admixture events that experience less recombination than those separated by larger genomic distances. A similar dip in $r_N$-$r_S$ values appeared in the *N. gonorrhoeae* data (Figure 2), although other evolutionary processes such as negative selection likely play a role. More complex models of selection will help further characterize these patterns.

**Balancing selection.**

Balancing selection may also produce local regions of high diversity and create coupling linkage between NSyn alleles if selection maintains multiple haplotypes at intermediate frequencies despite drift and recombination. To confirm this intuition, we simulated spatially variable selection (a type of balancing selection) in which two demes experience divergent selection pressures (Figure S14D), for example due to metabolic or immunological niches. Recombination between demes occurred frequently, and a genetic sample was taken from the metapopulation, randomly sampling from both demes in proportion to their size. As with simulations of adaptive admixture, balancing selection inflated NSyn couplings between close alleles (Figure S14E,F) and did so quite strongly when alleles under selection had intermediate effect sizes ($Ns$±10; Figure S14F). While simulations in Figure S14 were for equally sized demes, linkage dynamics were similar when demes were of unequal sizes (Figure S16).

As above, we also tested whether $r_N$ was significantly greater than $r_S$ using the Wilcoxon rank-sum test in two ways. Using the simulation results of balancing selection with effect sizes of $Ns$±4 (as shown in Figure 3D in the main text), we first applied this test to all common alleles (20-80% frequency) within 300bp and found significantly greater coupling linkage between neighboring NSyn alleles in 100% of simulation replicates, at the $\alpha$=0.05 significance level. We then further controlled for allele frequency differences between Syn and NSyn SNPs by binning common alleles into 10% frequency intervals, performing a significance test for each interval, and meta-analyzing all *P* values, as described above. For this analysis, 83% of simulation replicates had significantly more NSyn coupling linkage.

**Panmixia with positive or negative directional selection.**

To gain further intuition behind the behavior of $r_N$-$r_S$, we also simulated positive and negative directional selection within a single panmictic population under a variety of recombination rates. We first modeled selection with multiplicative effects (Figure S17), and none of these simulations produced positive $r_N$-$r_S$ between close alleles when calculated using all SNPs as in Figure 2A. Analysis of common alleles (20-80% frequency) within 300bp also showed $r_N$-$r_S$ was significantly positive (α=0.05) in only ~5% of replicates (Figure S17)

We next modeled directional selection with positive epistatic interactions between mutations, calculating individual fitness using the model of Desai *et al.* 2007 (Supplementary Methods). In the context of positive selection, positive fitness interactions correspond to synergistic epistasis between beneficial mutations. In the context of negative selection, they correspond to antagonistic epistasis between deleterious mutations. For both of these scenarios, we again find $r_N$-$r_S$ is not systematically positive for linked alleles when calculated using all SNPs, and analyses of common alleles within 300bp were only significant for ~5% of replicates (Figure S18)

#### SUPPLEMENTARY METHODS

##### Simulations of adaptive admixture.

Our model of admixture is distinct from traditional ones (e.g. Wright 1931) in that migration does not involve the movement of individuals or entire genomes between demes but only the transfer of small, geometrically distributed DNA fragments from a donor population (i.e. interspecies recombination).

We simulated 100kb fragments in a small sink population (or DNA recipient) of size $N_1$=2000 haploids and a larger source population (or DNA donor) of size $N_2$=10000. As shown in Figure S14A, these populations evolved in allopatry for $T_d=10N_1$ generations, after which unidirectional admixture from donor to recipient occurred for $T_a$ generations. In Figure S14B,C, $T_a$ was $0.1N_1$ generations but was varied from $0.02N_1$ to $2N_1$ generations in Figure S15. Throughout the entire simulation, neutral mutations arose within the metapopulation with rate $\theta=2(N_1+N_2)\mu=0.02$ per site (or $\theta_1=2N_1\mu=0.0033$ per site), where $\mu$ is the mutation rate per chromosome per generation. Each new mutation had a 1% chance of having a beneficial effect of size $s$, which was varied from $s=5\times10^{-5}$ to $s=1.25\times10^{-2}$ (or $N_1s=0.1$ to $N_1s=25$; Figure S15). Mutations that affected fitness had multiplicative effects, where individual fitness ($w$) was calculated as

$$w = (1+s)^m$$

where $m$ is the number of mutations with fitness effects. Recombination events arose twice as frequently as mutations with rate $\rho=2(N_1+N_2)r=0.04$ per site (or $\rho_1=2N_1r=0.0066$ per site), where $r$ is the mutation rate per chromosome per generation. Homologous recombination was modeled as gene conversion, and DNA tract lengths were geometrically distributed with a mean of 300bp. During $T_d$, recombination was only allowed between chromosomes from the same subpopulation, but during $T_a$ there was a 2% chance that DNA from the donor population served as template for recombination within the recipient population. For all parameter sets, we simulated 300 replicates.

After $T_d+T_a$ generations of evolution, we sampled $n=200$ chromosomes from the recipient population. We calculated $r_N$-$r_S$ as a function of distance between SNP pairs (Figure S14-S15) but also by "gene" (Figure 5), defined as 1kb segments within the simulated 100kb fragment. As noted in Results, we used a cutoff value of $r_N$-$r_S > 0.2$ because (1) it contained the top 10% of observed values in the UK genomic data and (2) this value was near the 95% quantile (0.23) of the distribution of intragenic $r_N$-$r_S$ from simulations with neutral effects (Figure 5).

#### Simulations of balancing selection.

To simulate spatially variable selection, a form of balancing selection, we evolved two subpopulations (or demes) in which mutations beneficial in one deme were deleterious in the other (Figure S14D). Migration between demes was frequent, with a 50% chance of DNA from deme 2 serving as template for recombination for chromosomes within deme 1, and vice versa. As above, neutral mutations arose within the metapopulation with rate $\theta=2(N_1+N_2)\mu=0.02$ per site, where subscripts indicate deme 1 or 2. However, each new mutation had a 5% chance of having a beneficial effect of size $s$, which was varied to give values of $N_1s=2$ or $N_1s=10$. Recombination events arose twice as frequently as mutations with rate $\rho=2(N_1+N_2)r=0.04$ per site. We simulated a metapopulation with equally sized demes of $N_1=N_2=1000$ (Figure S14E-F) or unequally sized demes of $N_1=200$ and $N_2=1800$ (Figure S16). For these simulations, we modeled shorter 10kb fragments, since patterns of $r_N$-$r_S$ for SNPs separated by distances longer than the mean recombination tract length (300bp) were similar.

**Simulations of panmixia with negative or positive directional selection.**

For simpler models of negative or positive directional selection, we simulated a 100kb segment from a single population of size $N$=10000 for 10$N$ generations with population mutation rate $\theta$=0.01 per site. We varied recombination rates, with $\rho/\theta$ = 0.1, 1, or 10, and the average length of DNA exchanged was 300bp, as above. For simulations of negative selection, mutations had a 66% chance of having a weak ($Ns$=-5) or strong ($Ns$=-50) deleterious effect, modeling a scenario in which mutations in the first two positions of a codon are nonsynonymous. For simulations of positive selection, mutations had a lower, 10% chance of having a weak ($Ns$=5) or strong ($Ns$=50) beneficial effect. As above, we calculated $r_N$-$r_S$ using a sample of $n$=200 chromosomes.

We first simulated selection with multiplicative fitness effects (Figure S17), where individual fitness was calculated as above. We next added positive epistatic interactions. Following Desai *et al.* 2007, we calculated individual fitness ($w$) with epistatic interactions as:

$$w = e^{sm^{1+\varepsilon}}$$

where $\varepsilon$ represents the sign and strength of epistasis. For beneficial mutations with positive values of $s$, positive values of $\varepsilon$ create synergistic epistasis, and combinations of mutations have a greater effect than the product of their individual effects. Likewise for deleterious mutations with negative values of $s$, negative values of $\varepsilon$ create antagonistic epistasis, with combinations of mutations having less of a deleterious effect than the product of their individual effects. For simulations of positive and negative selection with epistasis (Figure S18), we set $\varepsilon$ to 0.8 or -0.8, respectively. For other parameters, we used similar values to those for selection with multiplicative effects with the exception that we only specified weak individual effects for mutations; we did this for mutations under negative selection ($Ns$=-5), since stronger effects limited the number of common alleles, and also for mutations under positive selection ($Ns$=1), so that beneficial mutations persisted in the population long enough to interact with other mutations.

**Significance testing.**

As with the observed data, we tested if $r_N$-$r_S$ was significantly positive for common alleles (20-80%) in simulated data using the Wilcoxon rank-sum test ("Stats" package in R). When common alleles were further binned into 10% frequency intervals to compare alleles of more-similar frequency, we applied the Wilcoxon rank-sum test to each bin and meta-analyzed $P$ values across bins using Stouffers Z-score method ("metap" package in R; Liptak 1958). For selected parameter sets, we did these tests for each of

300 simulation replicates and reported the fraction of replicates in which $r_N$ was significantly greater than $r_S$ at the $\alpha=0.05$ significance level, indicating significantly more NSyn coupling linkage.

***dN/dS* analysis.**

To calculate values of *dN/dS* for each gene that also had high values of $r_N$-$r_S$ (Table S4), we used *omegaMap* with the following parameters: niter=30000, thinning=100, muPrior=improper_inverse, kappaPrior=improper_inverse, omegaPrior=inverse, indelPrior=improper_inverse, omega_model=constant, rho_model=constant, muStart=0.2, kappaStart=3.9, and indelStart=0.3. We also input codon frequencies estimated from the FA1090 reference sequence for *N. gonorrhoeae*. We discarded the first 20000 iterations of the MCMC chain and computed the mean value obtained from the last 10000 iterations. For each gene, this analysis was done for all three datasets, and we report the maximum observed *dN/dS* value in Table S4, although values across datasets were highly similar.

### SUPPLEMENTARY FIGURES

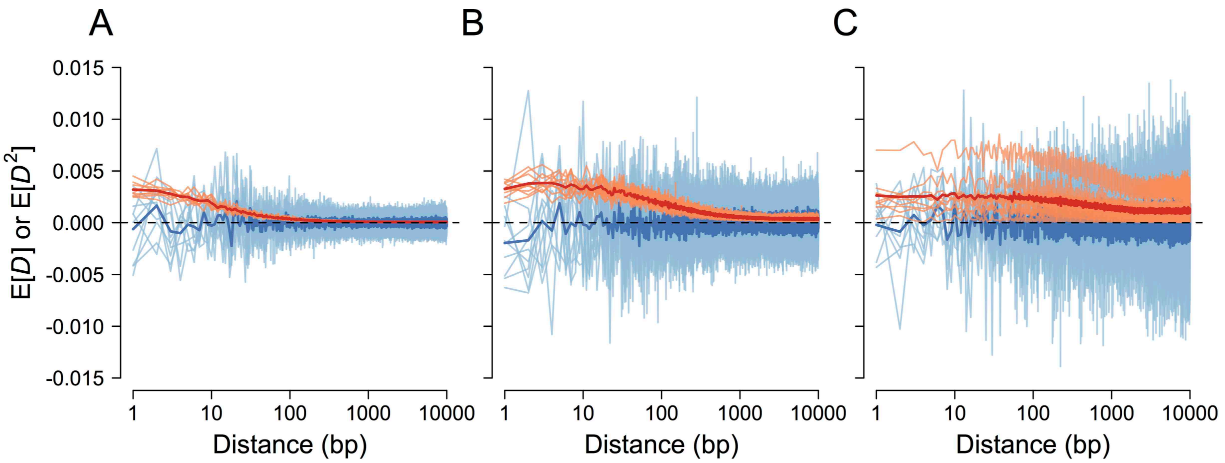

**Figure S1. E[D]=0 regardless of recombination.**
As previously shown by Hill and Robertson (1968) but confirmed here for a model of bacterial recombination, E[$D$] (blue lines) is zero for SNP pairs separated by any distance. However, E[$D^2$] (red lines), and thus E[$r^2$], becomes more positive for nearby SNPs due to limited recombination. Light blue or red lines represent expected values for individual simulations (10 in total), and dark blue or red lines show the overall mean. Here, we simulated 100kb fragments for $N$=10000 individuals with $\theta=2N\mu=0.01$ and $\rho/\theta$ = 10 (**A**), 1 (**B**), or 0.1 (**C**), where $\rho=2Nr$. Recombination tract lengths followed a geometric distribution with a mean of 1kb. For each simulation, we sampled $n$=200 individuals.

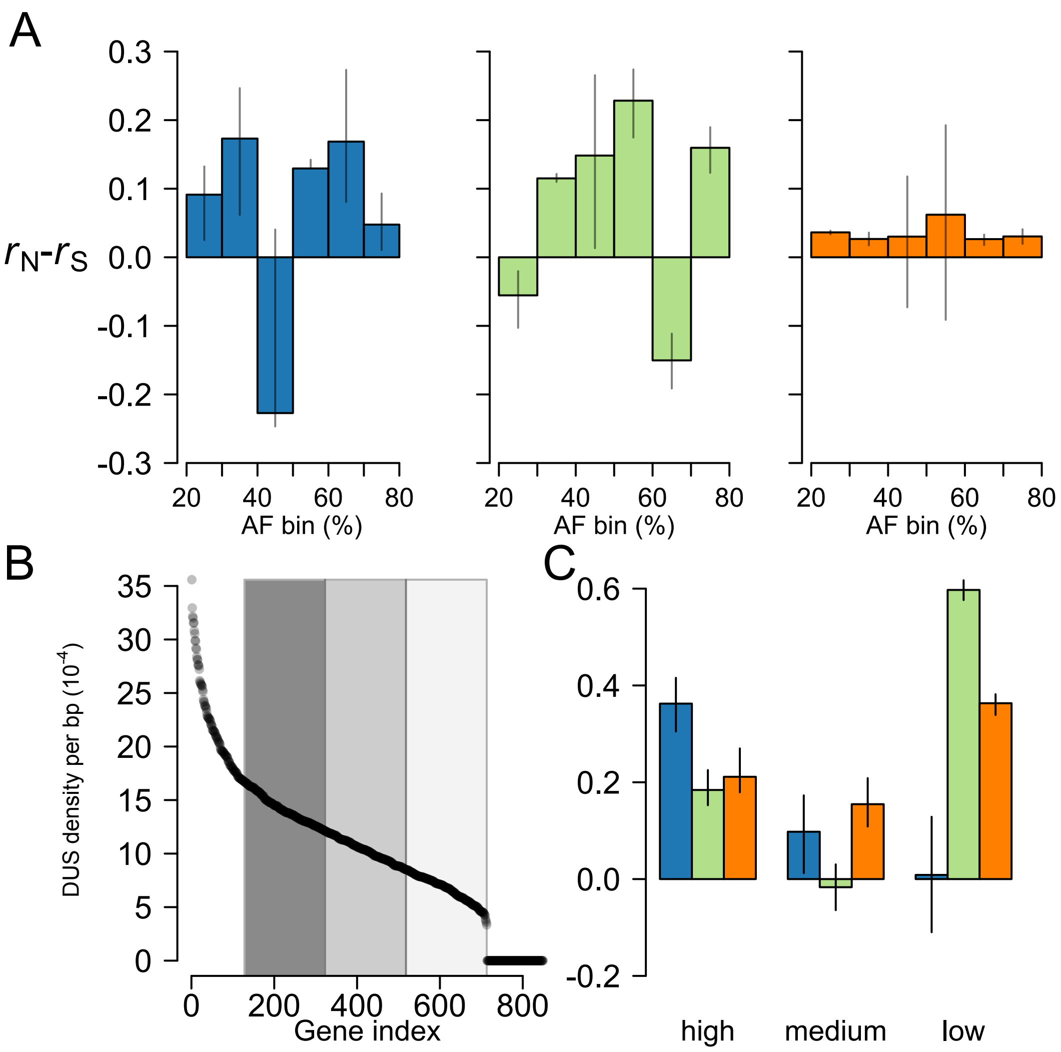

**Figure S2. Values of rN-rS for SNPs within 300bp while controlling for allele frequency and DUS density.**

**(A)** To control for potential allele frequency (AF) differences between Syn and NSyn SNPs, we binned intermediate-frequency alleles (20-80%) into 10% frequency intervals and calculated $r_N$-$r_S$ for each interval. **(B)** To control for relative differences in recombination frequency along the genome, we binned genes into three categories of DUS density: high (dark gray), medium (gray) and low (light gray). We excluded genes with no DUSs (right end of plot) and genes with the most DUSs (top 15% quantile; left end of plot) to avoid creating categories that contain genes with highly variable DUS densities. **(C)** Values of $r_N$-$r_S$ for genes within each DUS category. Analyses showing the overall significance of these positive deviations in $r_N$-$r_S$ can be found in Table S1, S2, and S3. Error bars in the barplots indicate the 5% and 95% quantiles of $r_N$-$r_S$ after subsampling the more frequent category of SNPs (*i.e.* Syn SNPs).

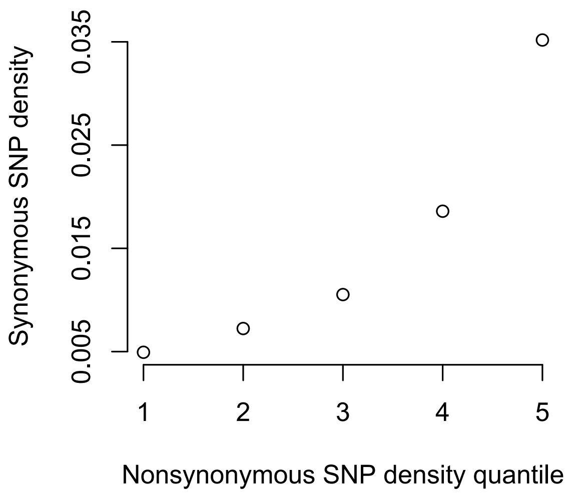

**Figure S3. Syn SNP density increases with NSyn SNP density.**
Levels of Syn SNP density (y-axis) for each NSyn SNP density quantile (x-axis) shows that genes with more NSyn SNPs tend to also have more Syn SNPs. These data are for the UK dataset.

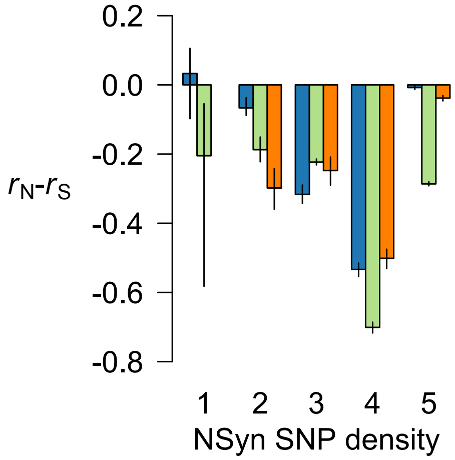

**Figure S4. NSyn linkage for rare variants.**
Shown are values of $r_N$-$r_S$ calculated only for rare variants below 5% frequency in the sample for UK (blue), NZ (green), and US (orange). Values for the lowest NSyn SNP density quantile for the US dataset are missing from too few NSyn SNPs. Error bars indicate the 5% and 95% quantiles of $r_N$-$r_S$ after subsampling the more frequent category of SNPs (*i.e.* Syn SNPs).

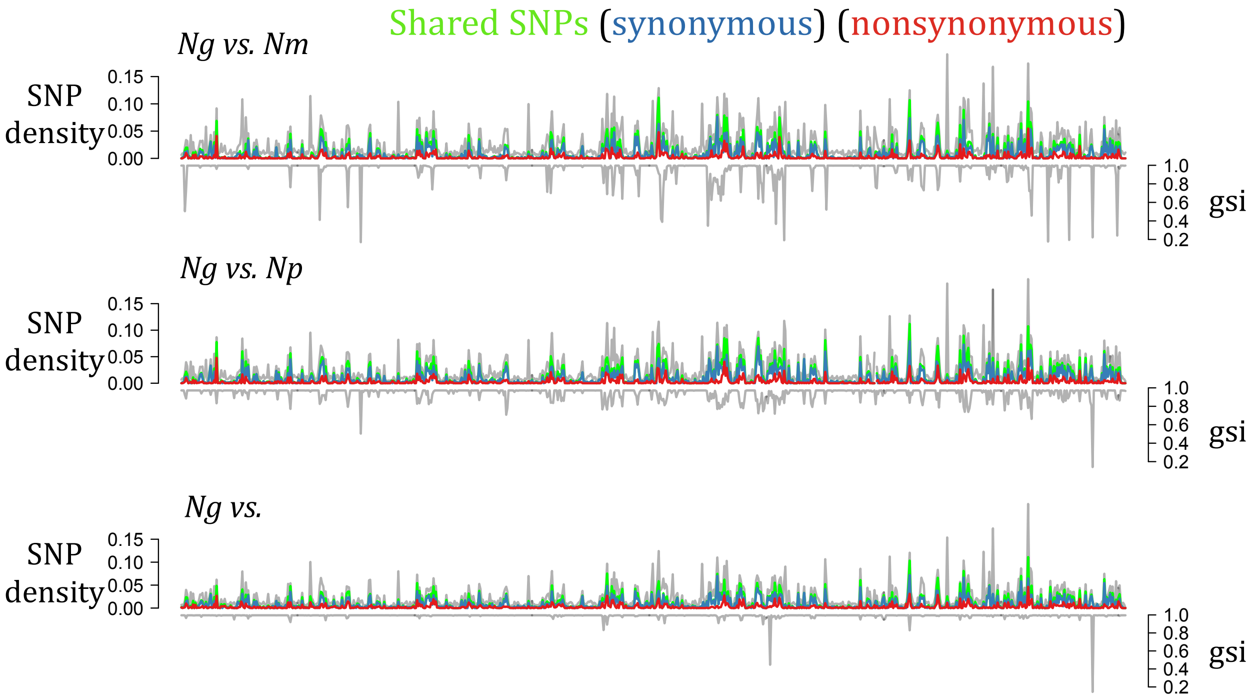

**Figure S5. Gene diversity correlated with interspecific-shared polymorphism and lower gsi values.** Diversity in *Ng* (only UK dataset shown), quantified with SNP density (gray lines in top panels), for each gene shows those with many SNPs also share many of these polymorphisms (green lines) with *Nm* (top), *Np* (middle), or *Nl* (bottom). Blue and red lines represent shared synonymous and nonsynonymous SNPs, respectively. These genes also have lower values of gsi (gray lines in bottom panels). Genes ordered in these plots according to their relative position in the FA1090 reference genome for *Ng*. The four *Neisseria* species used here are *N. gonorrhoeae* (*Ng*), *N. meningitidis* (*Nm*), *N. polysaccharea* (*Np*), and *N. lactamica* (*Nl*).

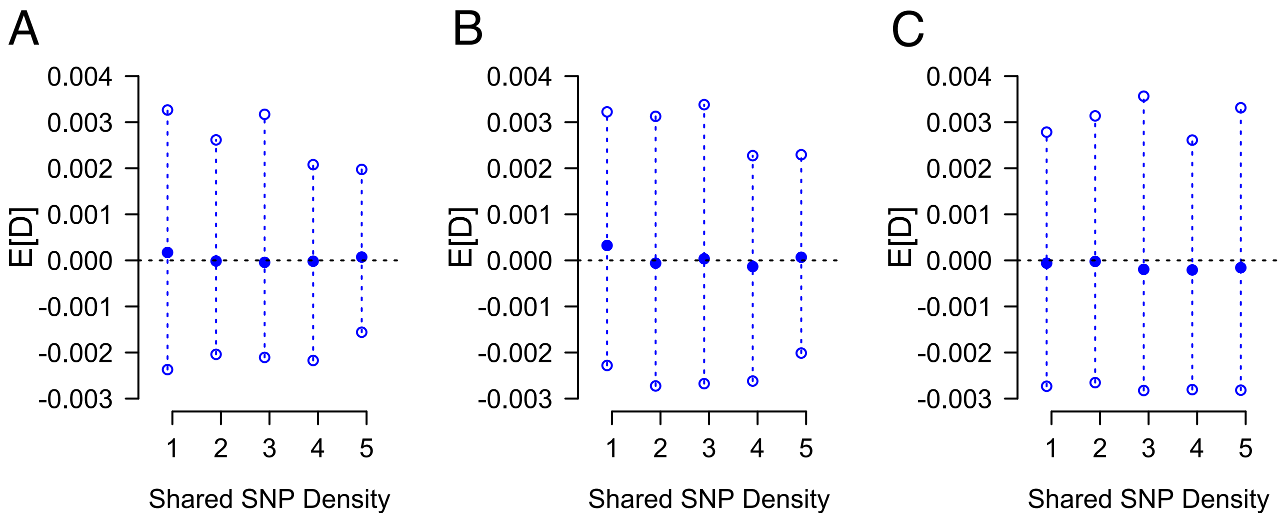

**Figure S6. Simulation confirm ILS does not inflate D***.*
We simulated a single population of 500kb DNA segments that split into two and took a sample (n=200) from one of these populations after various divergence times: 0.5*N* (A), 1*N* (B), and 2*N* (C) generations. We defined genes as 1kb regions within the 500kb segment and categorized these genes by the amount of SNPs that are also present in the second, unsampled population (i.e. shared SNP density). Since there is no admixture or gene flow between diverging populations, all shared SNPs must be due to incomplete lineage sorting (ILS), such that genes with higher shared SNP densities have deeper genealogies due to ILS. We then calculated mean values of *D*, or E[*D*], across all genes in a shared SNP density quantile to see if *D* became more positive with ILS. For samples taken after a variety of different divergence times, which should exhibit varying levels of ILS, mean values of *D* were not systematically more positive for genes with more evidence of ILS. Closed circles represent the median value of E[*D*] across 300 replicate simulations, and open circles represent the 5% and 95% quantiles. For these simulations, we set $\theta$=0.0043, $\rho$=0.0086, and the mean recombination tract length as 2500bp, as previously estimated for *N. gonorrhoeae* (Arnold *et al.* 2018).

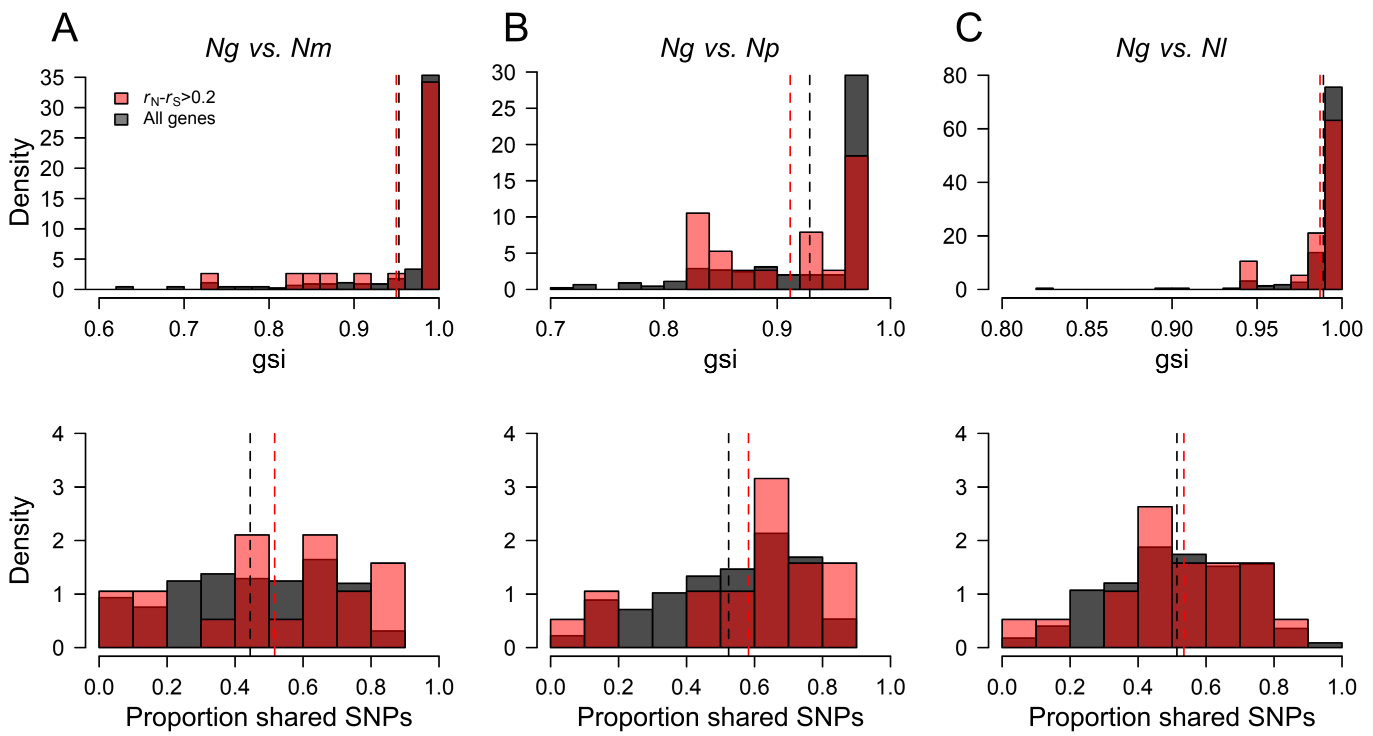

**Figure S7. Shared ancestry for genes with an excess of NSyn couplings.**
Histograms of gsi (top row) and shared SNP density (bottom row) are shown for all genes (black) or only those with $r_N$-$r_S$>0.2 (red) when comparing *N. gonorrhoeae* (*Ng*) with *N. meningitidis* (*Nm*; **A**), *N. polysaccharea* (*Np*; **B**) and *N. lactamica* (*Nl*; **C**). Darker red areas show where histograms overlap, and dashed lines show overall means.

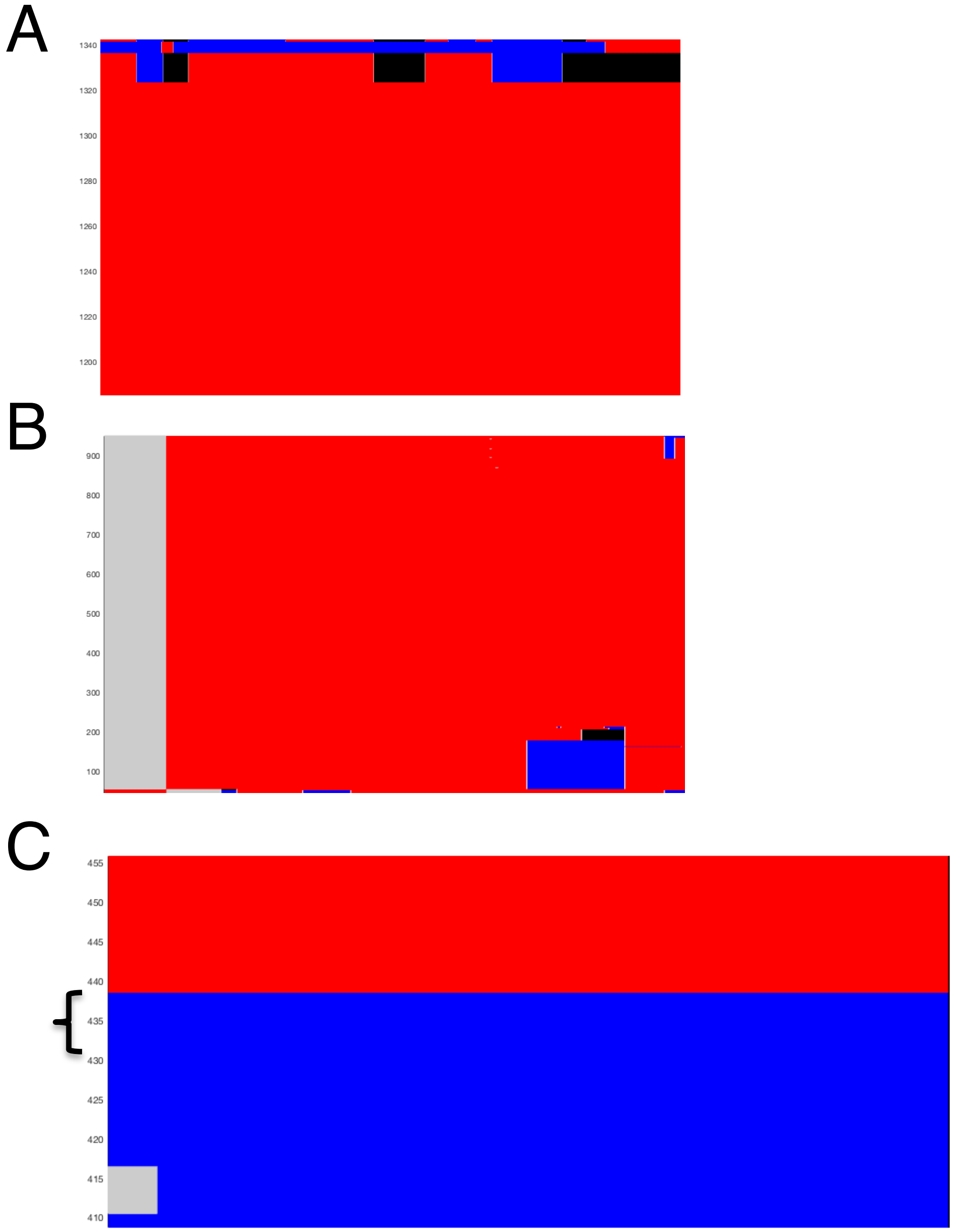

**Figure S8. Three examples of high-resolution fastGEAR admixture plots of genes with high $r_N$-$r_S$.** (**A**) When analyzing NGO0820 alleles (1,311 bp) from *N. gonorrhoeae* (red) along with *N. meningitidis*, fastGEAR detects DNA tracts that likely come *N. meningitidis* (blue) or another unknown species (black). (**B**) Likewise, when analyzing NGO0827 alleles (618 bp) from *N. gonorrhoeae* (red) along with *N. polysaccharea*, fastGEAR identifies donor DNA from *N. polysaccharea* (blue) along with other DNA from an unknown species (black). Part (**A**) is zoomed in and only shows a subset of the total dataset to better visualize the admixed alleles. Although we included multiple species in **A** and **B**, only *N. gonorrhoeae* alleles are shown here. In (**C**), both *N. gonorrhoeae* (red) and *N. meningitidis* (blue) alleles for NGO1602 are shown, but those to right of the black curly bracket came from *N. gonorrhoeae* isolates. Gray regions indicate gaps in the alignment. Nucleotide position is represented along the x-axis, and alleles are represented as horizontal color bars and are indexed numerically along the y-axis. All of these results agree with values of gsi for both genes in Table S4.

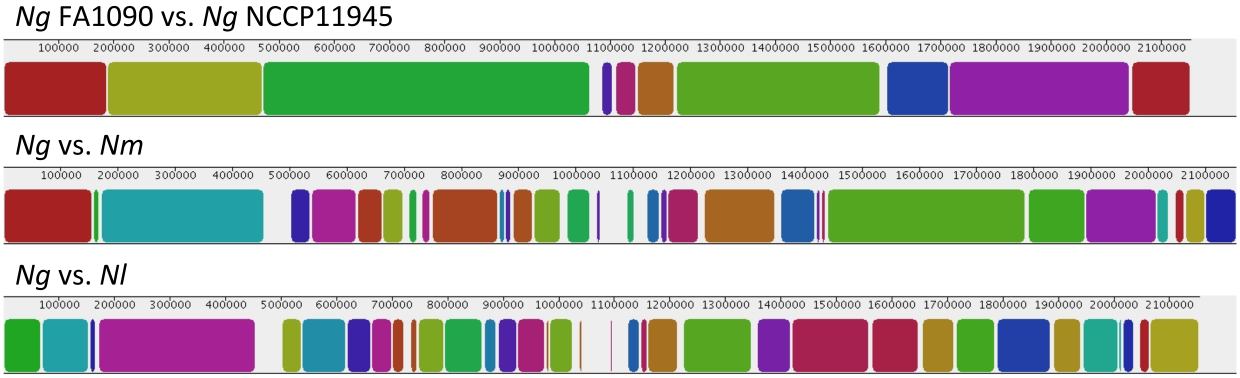

**Figure S9. Microsynteny between *Neisseria* used in this study.**
We used progressiveMauve to align the *N. gonorrhoeae* (*Ng*) FA1090 reference genome to another *Ng* reference NCCP11945 but also to the closely related species *N. meningitidis* (*Nm*) and *N. lactamica* (*Nl*). Colored blocks represent the length of syntenic bocks identified between the two genomes listed. While these data provide evidence of rearrangements within *Ng*, syntenic blocks generally extend distances much longer than 30kb, the maximum distance used in Figure1 and Figure 2. However, genomic distances greater than 300kb may be less accurate. *N. polysaccharea* was not included because there is currently no completed reference genome for this species.

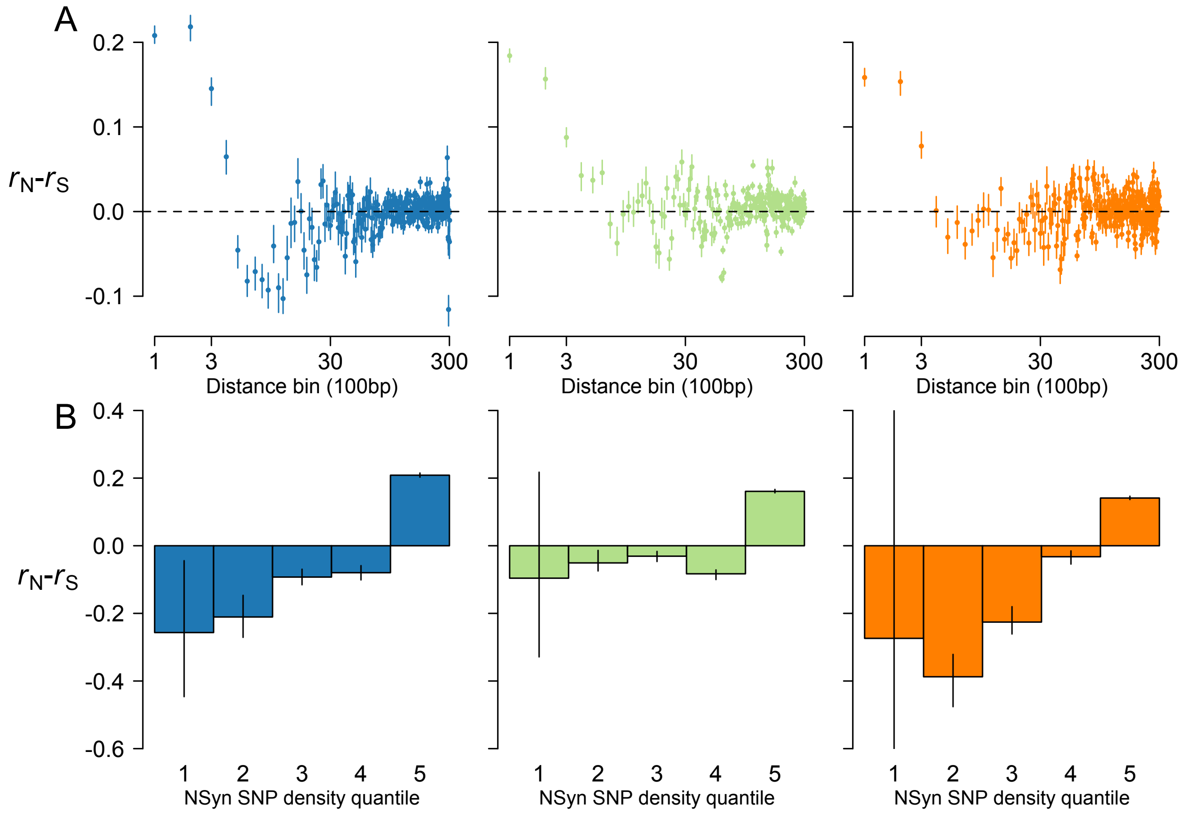

**Figure S10. Calculating $r_N$-$r_S$ with additional sets of outgroup sequences.**
Here we calculate $r_N$-$r_S$ but polarize mutations by only considering sites that are identical and invariant in both the *N. meningitidis* and *N. polysaccharea* genome. Alleles at these sites in the outgroups were considered the ancestral state for a biallelic SNP in *N. gonorrhoeae*. **(A)** $r_N$-$r_S$ for SNP pairs within the same distance bin. **(B)** $r_N$-$r_S$ within genes, categorized by their NSyn SNP density. Results are shown for UK (blue), NZ (green), and US (orange). Error bars indicate the 5% and 95% quantiles of $r_N$-$r_S$ after subsampling the more frequent category of SNPs (*i.e.* Syn SNPs).

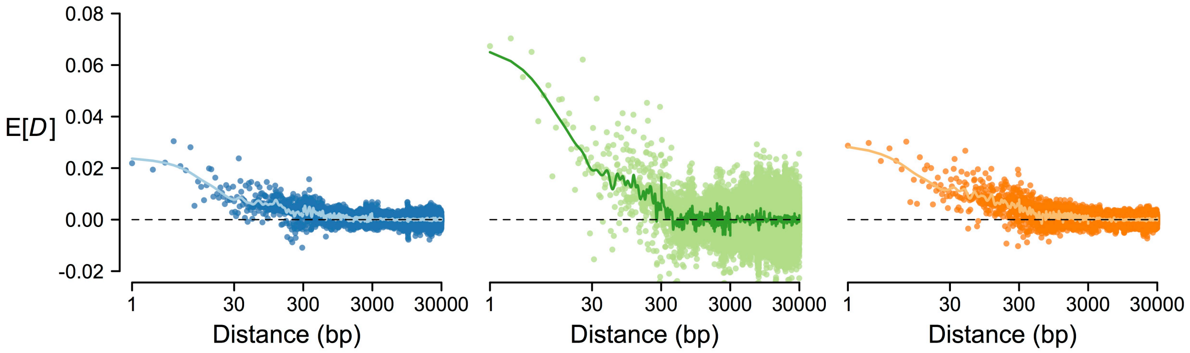

**Figure S11. Patterns of *D* when only excluding identical isolates.**
Shown are values of E[*D*] as a function of distance, as in Figure 1B, except values were calculated from larger datasets in which only identical isolates were removed, compared to analyses in the main text in which isolated with fewer than 6 SNP differences were removed.

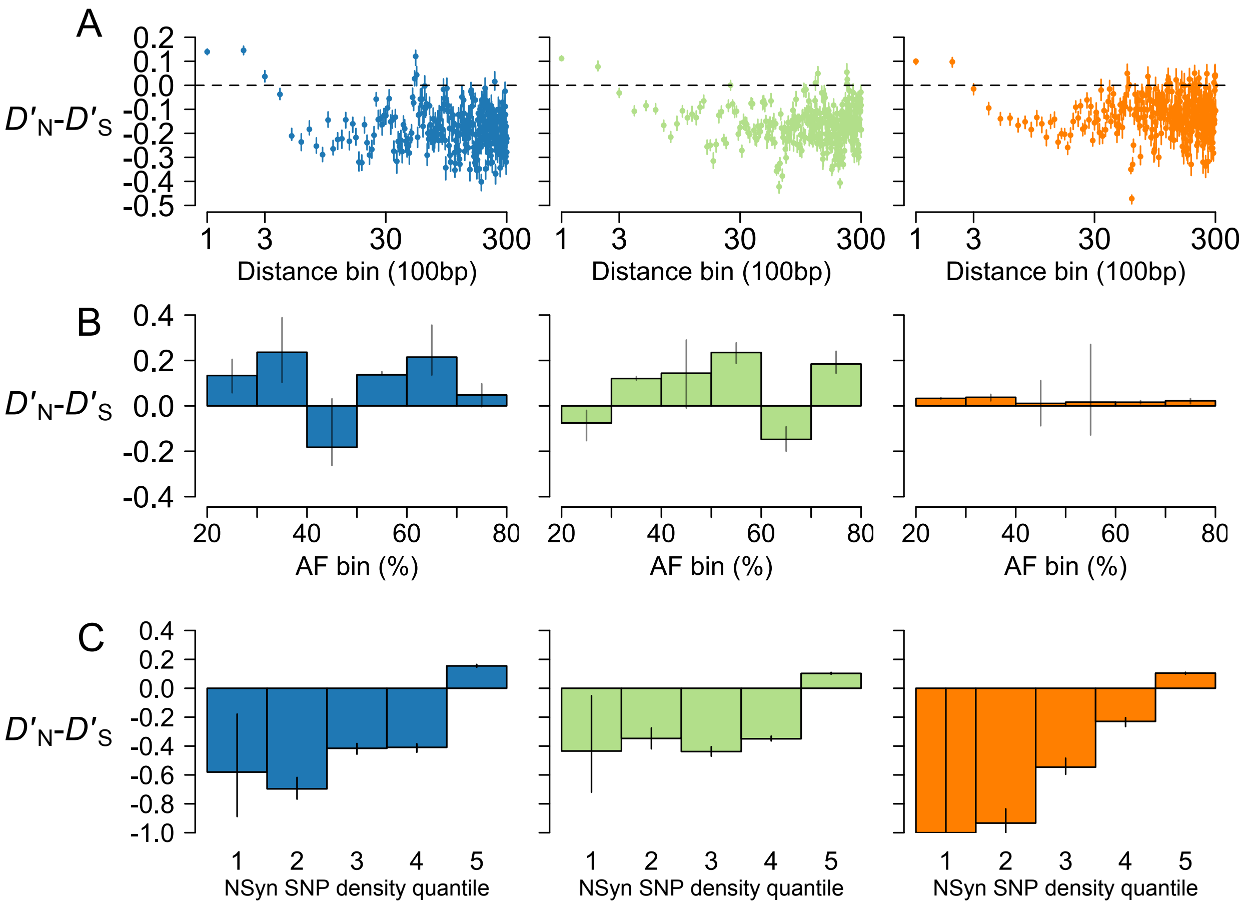

**Figure S12. Reanalysis of NSyn and Syn pairwise associations with D'.**
**(A)** We binned SNP pairs by distance (100bp intervals) and compared NSyn *D'* ($D'_N$) to Syn *D'* ($D'_S$) for pairs within the same distance bin. **(B)** Focusing on SNPs less than 300bp apart, we calculated $D'_N$-$D'_S$ only for intermediate-frequency alleles (20-80%), binning alleles into 10% frequency intervals as in Figure S2. **(C)** We then categorized genes by NSyn SNP density and calculated $D'_N$-$D'_S$ within genes. $D'_N$-$D'_S$ was significantly greater than zero only for genes with many NSyn SNPs. Results are shown for UK (blue), NZ (green), and US (orange). Error bars indicate the 5% and 95% quantiles of $r_N$-$r_S$ after subsampling the more frequent category of SNPs (*i.e.* Syn SNPs).

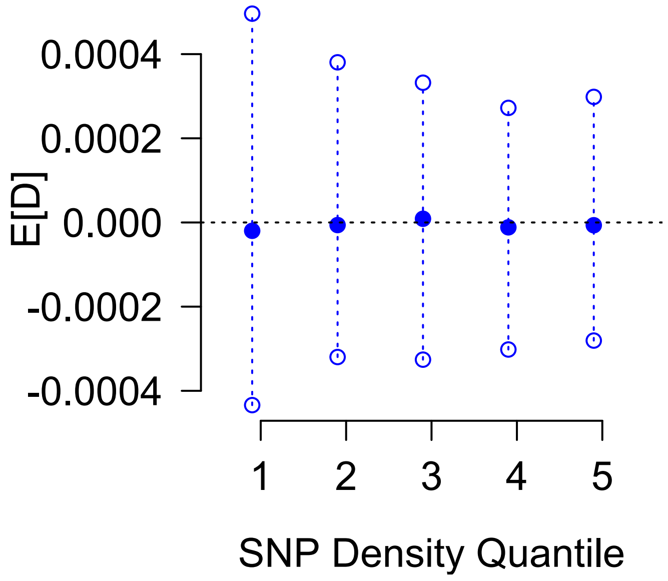

**Figure S13. Mean value of *D* does not change with SNP density.**
We simulated 500kb segments under neutrality, specifying each 1kb interval as a "gene", and categorized these genes by SNP density. We then calculated mean values of *D* for all genes in a SNP density quantile for each of 300 simulations. Shown are the median values of E[*D*] (closed dots) and the 5% and 95% quantiles (open dots). For each simulation, $\theta$=0.0043/bp, $\rho$=0.0086/bp with a mean tract length of 2.5kb (as previously estimated for *N. gonorrhoeae*; (Arnold *et al.* 2018), and we sampled n=200 haploid individuals to calculate summary statistics.

### SUPPLEMENTARY FIGURES FROM SIMULATIONS

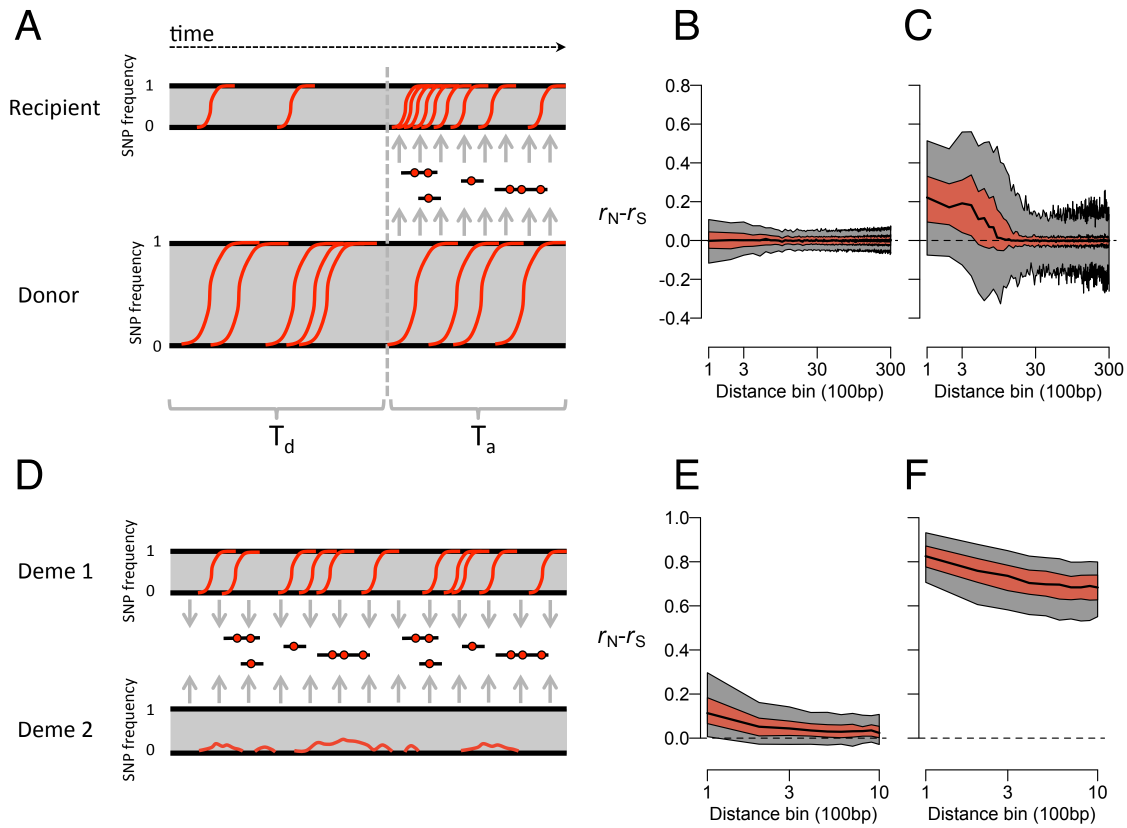

**Figure S14. Simulating the effects of adaptive admixture and balancing selection on $r_N$-$r_S$.**
(**A**) We simulated two allopatric populations, represented here as SNP frequencies through time. For $T_d$ generations, beneficial mutations randomly arise and may increase in frequency (red trajectories). Afterwards, unidirectional admixture from the donor to the recipient population occurs over $T_a$ generations, during which the recipient population receives linked, beneficial mutations (red dots). At time $T_d + T_a$, we sample sequences from the recipient population and calculate $r_N$-$r_S$. Neutral mutations that randomly arise much more frequently than beneficial ones are not shown. When beneficial mutations have nearly neutral effect sizes (*Ns*=0.1;**B**), $r_N$-$r_S$ is ~0 for neighboring or distant SNPs but becomes positive for neighboring SNPs once beneficial effect sizes become intermediate in strength (*Ns*=25;**C**). The central black line represents the median $r_N$-$r_S$, the red region spans the interquartile range of medians, and the gray regions spans the 5% to 95% quantile of medians across 300 replicates. For these simulations (**B,C**), we model 100kb fragments with a recombination tract length of 300bp. **(D)** We also simulated spatially variable selection where mutations are beneficial in deme 1 but deleterious in deme 2, with demes being equal in size. $r_N$-$r_S$ is slightly greater than zero for neighboring SNPs with weak selection (*Ns*=2; **E**) but much greater than zero for all SNPs with intermediate selection (*Ns*=10; **F**). For these simulations (**E,F**) we model 10kb fragments with the same recombination parameters as **B**,**C**.

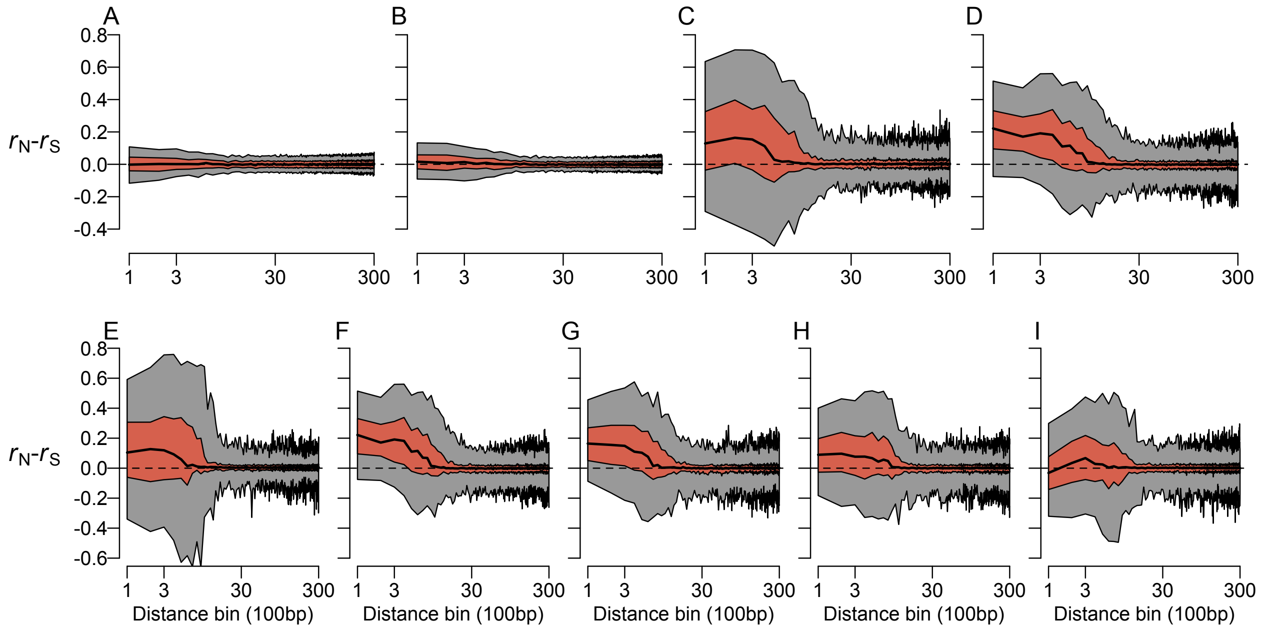

**Figure S15. Behavior of $r_N$-$r_S$ for simulations of adaptive admixture across a range of parameter space.**

When beneficial mutations had effect sizes that were nearly neutral (*Ns*=0.1; **A**) or weak (*Ns*=1; **B**), $r_N$-$r_S$ was roughly 0 for neighboring or distant SNPs. However, $r_N$-$r_S$ tended to be positive for nearby SNPs once beneficial effect sizes became intermediate in strength (*Ns*=10 or 25; **C** and **D**, respectively). For these simulations (**A**-**D**), samples were taken from the recipient populations 0.1*N* generations after the onset of admixture. The central black line represents the median $r_N$-$r_S$, the red region spans the interquartile range of medians, and the gray regions spans the 5% to 95% quantile of medians across 300 replicate simulations. Panels **E-I** show the effect on $r_N$-$r_S$ of sampling the recipient population after the onset of admixture (with *Ns*=25) at different times: 0.02*N* (**E**), 0.1*N* (**F**), 0.5*N* (**G**), 1*N* (**H**), or 2*N* (**I**) generations.

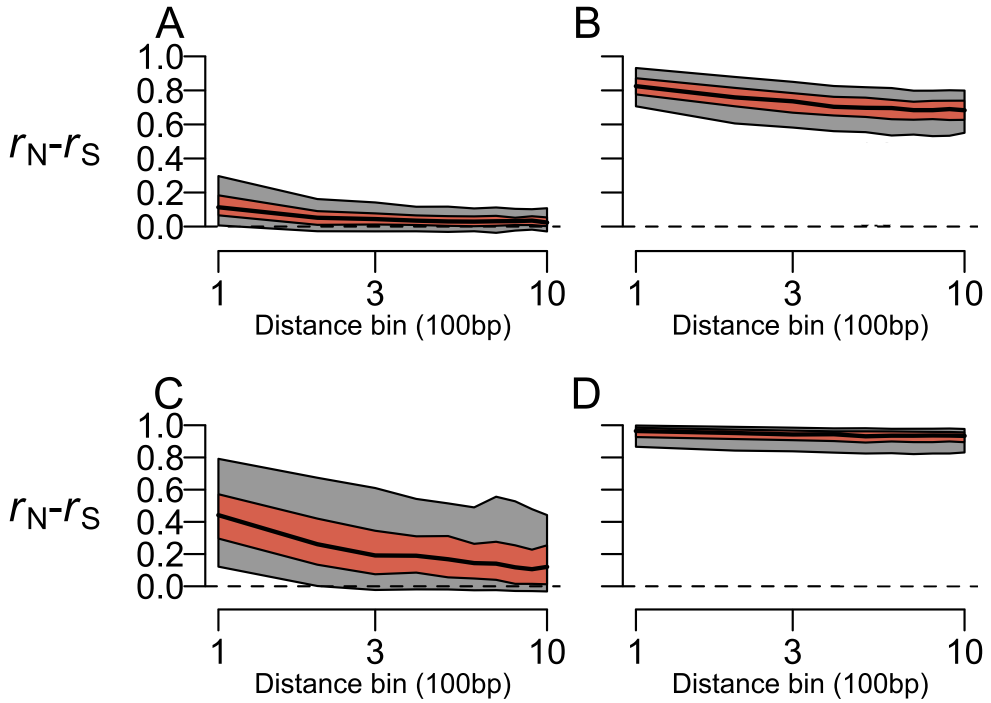

**Figure S16. Balancing selection with demes of equal and unequal sizes.**
Shown are simulations of spatially variable selection when subpopulations are of equal size (*N*1=*N*2=1000; **A,B**) or when one deme is 10% of the size of the total population (*N*1=200, *N*2=1800; **C,D**). The strength of selection also varies from weak (*Ns*=2; **A,C**) to intermediate (*Ns*=10; **B,D**).

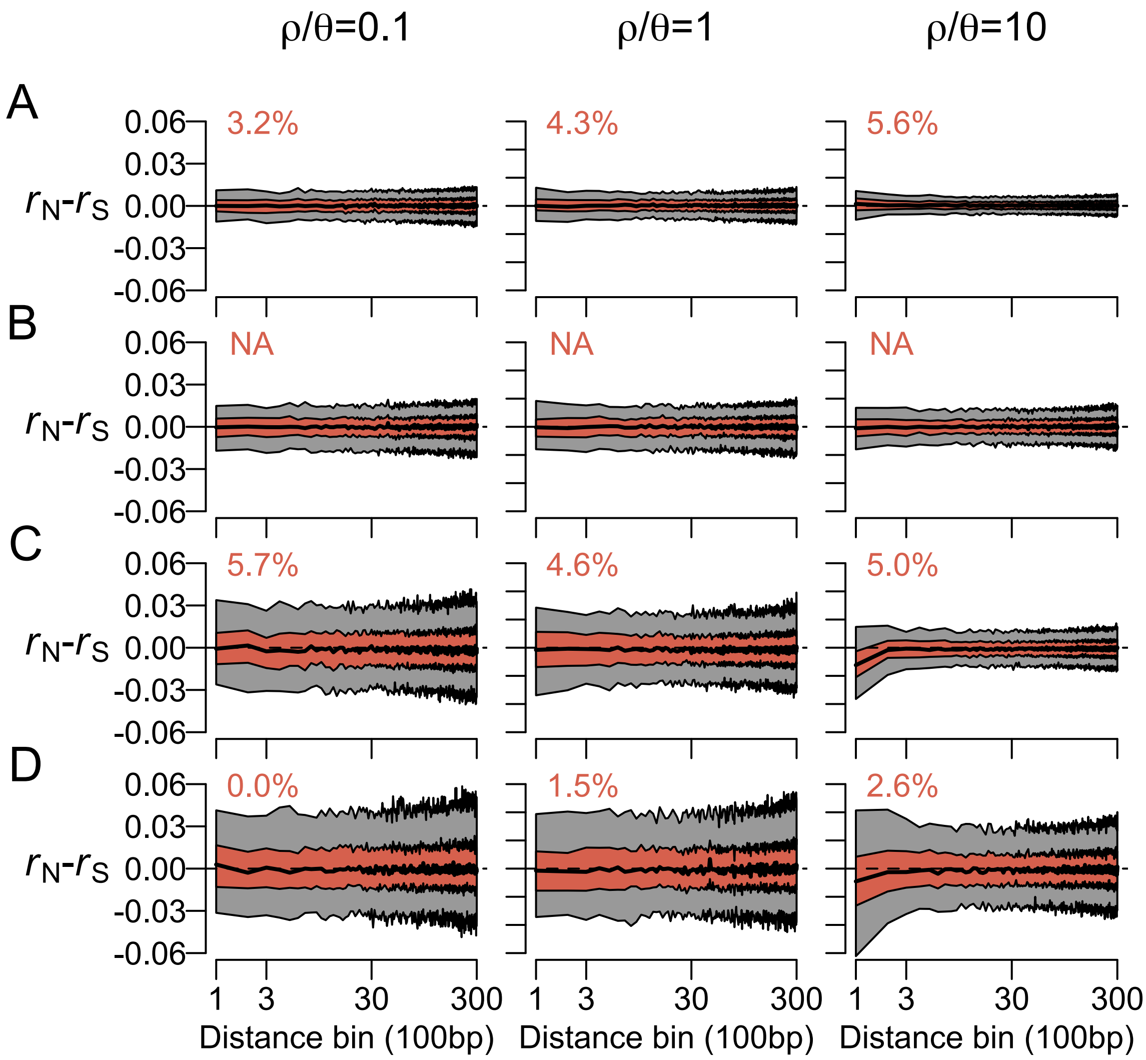

**Figure S17. Simulations of panmixia with negative or positive directional selection and multiplicative fitness effects.**
We simulated a single population under negative (**A,B**) or positive (**C,D**) selection. We explored a wide range of parameter space, including selective effects that were either weak (*Ns* = -5 in **A**; *Ns* = 5 in **C**) or strong (*Ns* = -50 in **B**; *Ns* = 50 in **D**) across a range of recombination rates, with $\rho/\theta$ =0.1, 1, or 10 for the left, middle, and right column, respectively. The percentages in red indicate the fraction of simulation replicates, for a particular set of parameters, in which $r_N$-$r_S$ was significantly positive for common alleles (20-80% frequency) within 300bp. For simulations of strong negative selection (**B**), we do not report significance because too few replicates had NSyn alleles above 20% frequency.

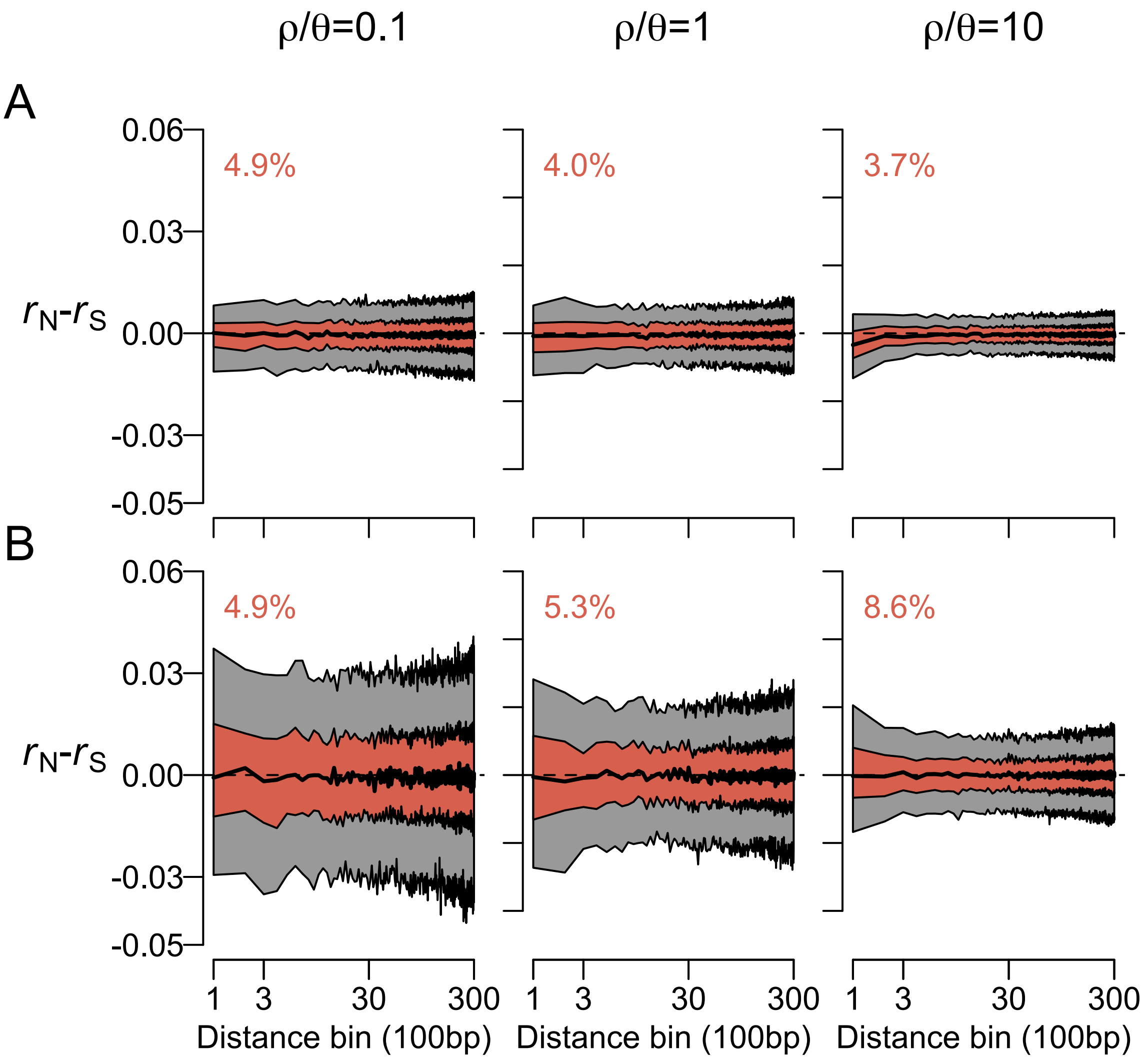

**Figure S18. Simulations of panmixia with negative or positive directional selection and epistatic fitness effects.**

We simulated a single population under negative (**A**) or positive (**B**) selection with epistatic fitness effects. We specified weak individual effects of mutations ($Ns$ = -5 in **A**; $Ns$ = 1 in **B**), but these effects interact in a positive way ($\varepsilon$ = -0.8 for antagonistic epistasis in **A**; $\varepsilon$ = 0.8 for synergistic epistasis in **B**). We explored these scenarios across a range of recombination rates, with $\rho/\theta$ =0.1, 1, or 10 for the left, middle, and right column, respectively. The percentages in red indicate the fraction of simulation replicates, for a particular set of parameters, in which $r_N$-$r_S$ was significantly positive for common alleles (20-80% frequency) within 300bp.
